# Amyloid Precursor Protein (APP) controls excitatory/inhibitory synaptic inputs by regulating the transcriptional activator Neuronal PAS Domain Protein 4 (NPAS4)

**DOI:** 10.1101/504340

**Authors:** Rémi Opsomer, Sabrina Contino, Florian Perrin, Bernadette Tasiaux, Pierre Doyen, Maxime Vergouts, Céline Vrancx, Anna Doshina, Nathalie Pierrot, Jean-Noël Octave, Serena Stanga, Pascal Kienlen-Campard

**Affiliations:** CEMO-Alzheimer Dementia group, Institute of Neuroscience, Université catholique de Louvain, Brussels, Belgium.; de Duve Institute, Ludwig Institute for Cancer Research and Université catholique de Louvain, Brussels, Belgium.; CEMO-Laboratory of Neuropharmacology, Institute of Neuroscience, Université catholique de Louvain, Brussels, Belgium.

**Keywords:** Alzheimer’s disease, APP family proteins, Neuronal differentiation, Transcriptome analysis, NPAS4, CRISPR-Cas9, Inhibitory neurotransmission.

## Abstract

Sequential proteolysis of the amyloid precursor protein (APP) and amyloid-β peptide (Aβ) release is an upstream event in Alzheimer’s disease (AD) pathogenesis. The function of APP in neuronal physiology is still, however, poorly understood. Along with its paralog APP-like Proteins 1 and 2 (APLP1-2), APP is involved in neurite formation and synaptic function by mechanisms that are not elucidated. APP is a single-pass transmembrane protein expressed at high levels in the brain that resembles a cell adhesion molecule or a membrane receptor, suggesting that its function relies on cell interaction processes and/or activation of intracellular pathways of signal transduction. Along this line, the APP intracellular domain (AICD) was reported to act as a transcriptional factor for targeted gene activation that mediates physiological APP functions. Here, we used an unbiased transcriptome-based approach to identify the genes transcriptionally regulated by APP in the rodent embryonic cortex and upon maturation of primary cortical neurons. The transcriptome analysis did not detect any significant differences in expression of previously proposed AICD target genes. The overall transcriptional changes were subtle, but we found that genes clustered in neuronal-activity dependent pathways are dysregulated in the absence of APP. Among these genes, we found the activity-dependent Neuronal PAS domain protein 4 (NPAS4) Immediate Early Gene to be downregulated in the absence of APP. Down-regulation of NPAS4 in APP knock-out (KO) neurons is not related to AICD but to the APP ectodomain. We studied the effect of APP deficiency on GABAergic and glutamatergic transmission, and found an increased production of the inhibitory neurotransmitter GABA in APP KO neurons, along with a reduced expression of the GABA(A) receptors alpha1, suggesting an impaired GABAergic neurotransmission in the absence of APP. CRISPR-Cas-mediated silencing of NPAS4 in neurons led to similar observations. Altogether, our results point out a new role for APP in the regulation of excitatory/inhibitory neurotransmission through the regulation of the activity-dependent NPAS4 gene.

## Introduction

The Amyloid Precursor Protein (APP) has been extensively studied as the precursor of the amyloid-β peptide (Aβ), the major component of the senile plaques which are a typical hallmark of Alzheimer’s disease (AD). Still, the physiological functions of APP *per se* have been largely overlooked and remain a matter of controversy. Understanding the physiological function of APP and how its deregulation would contribute to AD pathogenesis is thus of prime interest.

APP is a type 1 transmembrane protein that belongs to the APP-like protein family (APLP1 and APLP2, referred to as APLPs), which are present in most of the species, excepted in yeast, prokaryotes and plants. The APLP family has been generated by several duplications and contraction events during evolution. The specific physiological role and/or redundant functions assigned to each member are yet not clearly defined (for a review see Shariati and De Strooper, 2013). APP-/- mice show a subtle phenotype, with reduced body and brain weight, reduced locomotor activity, gliosis, mild axonal growth/white matter defects and altered long-term potentiation responses (Guo et al., 2012; Muller et al., 2012; Muller and Zheng, 2012). In this broad (and complex) picture, growing evidence indicate that APP controls neuronal proliferation, differentiation (Freude et al., 2011; Hu et al., 2013) and migration during embryogenesis (Young-Pearse et al., 2007). APP contributes to the establishment of a functional neuronal network by promoting neurite outgrowth (Hoe et al., 2009b). Additionally, APP was reported to control synaptic formation and activity (Priller et al., 2006; Santos et al., 2009; Lee et al., 2010; Pierrot et al., 2013; Klevanski et al., 2015; Zou et al., 2016) in the central nervous system (CNS) and at the neuromuscular junction (Stanga et al., 2016). APP directly modulates the excitatory neurotransmission by interacting with AMPA (Lee et al., 2010) or NMDA (Cousins et al., 2009; Hoe et al., 2009a) receptors. APP was also described to play an important role in GABAergic inhibitory neurotransmission. APP deficiency reduces paired pulse depression (PPD) in mice (Seabrook et al., 1999) and affects GABA receptors expressions (Fitzjohn et al., 2000; Chen et al., 2017), while APP overexpression induces hyperexcitability due to GABAergic neurotransmission failure (Born et al., 2014). Recently, APP was associated with the GABA excitatory/inhibitory shift occurring in embryonic neurons (Doshina et al., 2017). APP appears therefore to have a direct role in the fine-tuning of excitatory and inhibitory neurotransmission, a process that seems to be also critical in AD pathogenesis.

Tuning inhibitory/excitatory neurotransmission is very important for neuronal plasticity and memory formation. This is regulated by a specific subset of genes induced by neuronal activity, belonging to the Immediate Early Genes (IEGs) family, which control the mechanisms that “reshape” synaptic inputs on neurons (West and Greenberg, 2011). IEGs expression is instrumental to neuronal plasticity and memory formation (Alberini, 2009; Loebrich and Nedivi, 2009; Leslie and Nedivi, 2011). Among these IEGs, NPAS4 is specifically involved in a transcriptional program that regulates neuronal firing responses to excitatory transmission by enhancing inhibition (Lin et al., 2008), therefore keeping neuronal firing in response to stimuli within normal levels (Spiegel et al., 2014). Elevated activity of inhibitory neurons also induces NPAS4, promoting increased excitation onto the same neurons (Spiegel et al., 2014). NPAS4 is therefore a key player in the maintenance of excitatory/inhibitory balance in neuronal network.

The precise mechanisms underlying APP synaptic functions are still elusive. One could suspect APP to regulate the expression of genes involved in synaptic activity, or to shape the structure of the synapse. APP was shown to control gene expression through its intracellular domain called AICD. An increasing list of AICD candidate genes has emerged from various models (reviewed in Pardossi-Piquard and Checler, 2012). Some of these candidate genes failed to be confirmed by transcription analysis in APP-deficient cell lines (Hebert et al., 2006; Waldron et al., 2008), and APP was also reported to regulate gene transcription independently of AICD release (Hicks et al., 2013; Pierrot et al., 2013). It is so far impossible to clearly define (i) the precise identity of APP target genes in neurons (ii) how these APP target genes relate to APP neuronal function (iii) the mechanism involved in APP-dependent in gene transcription.

In the present study, we first aimed at identifying genes transcriptionally regulated by APP in primary neurons. To that end, we performed a non-biased transcriptome analysis of APP+/+ and APP-/- primary cortical neurons at different stage of differentiation. In-depth transcriptome analysis revealed that the absence of APP induced only subtle changes in global gene expression. The hitherto described AICD target genes were not significantly up-or down-regulated in our model. A more detailed analysis indicates that expression of genes clustered in specific neuronal pathways was affected by the absence of APP. In particular, the transcription of the activity-dependent transcription factor *Npas4* gene was down-regulated in the absence of APP after 7 days of culture. Interestingly, we observed that the amount of the inhibitory neurotransmitter γ-aminobutyric acid (GABA) and the expression of glutamate decarboxylase 65 (GAD65), the enzyme that catalyzes the decarboxylation of <u>glutamate</u> to <u>GABA</u>, were increased in APP-/- neurons, suggesting that the inhibitory inputs in synaptic transmission are increased in APP KO neurons. Direct down-regulation of *Npas4* by CRISPR-Cas9 editing in neurons mimicked the increase in GAD65 and GABA release observed in APP-/- cultures. Altogether, our data give a new in APP-dependent neuronal activity, supporting that APP tunes the excitatory/inhibitory transmission in neuronal networks.

## Materials and Methods

### Antibodies, chemicals and reagents

All media and reagents used for cell cultures were purchased from Thermo Fisher Scientific (Waltham, MA); fetal bovine serum was purchased from Biowest (Nuaillé, France). Analytical grade solvents and salts were purchased from Sigma-Aldrich (St-Louis, MO). N-[N-(3,5- Difluorophenacetyl)-L-alanyl]-S-phenylglycine t-butyl ester (DAPT), sAPPα (S9564) and DAPI (D9542) were from Sigma-Aldrich (St-Louis, MO, USA). Triton-X100 was purchased from Merck (Darmstadt, Germany) and TriPure Isolation Reagent from Roche (Basel, Switzerland). Microarray analysis kits were from Affymetrix (Santa Clara, CA, USA). All reagents for RNA processing or cDNA synthesis were purchased from Bio-Rad (Hercules, CA). Primers were purchased from Sigma-Aldrich (St Louis, MO, USA). Proteins were quantified with BCA Protein Assay kit (Thermo Fisher Scientific, Waltham, MA). NuPAGE® reagents were from Invitrogen (Carlsbad, CA). PVDF and nitrocellulose membranes were from Merck Millipore (Billerica, MA,) or Amersham™ (Little Chalfont, UK). Nonfat dry milk was from Merck (Darmstadt, Germany). Western Lighting® Plus-ECL reagents were from PerkinElmer (Waltham, MA) and Fluoprep mounting medium was from bioMérieux (Marcy l’Etoile, France). Lentivirus were prepared with Acrodisc® 0,45µm filters (Pall, NYC, USA) and LentiX™ Concentrator reagent (Clontech, Mountain View, CA). The following antibodies were used: APP NT (22C11, MAB348, Merck Millipore, Billerica, MA), anti-human APP (WO2, MABN10, Merck Millipore, Billerica, MA), anti-APP CT (Y188, Abcam, Cambridge, UK), anti-APLP1 (Cat. No. 171615, Calbiochem EMD Biosciences – Merck, Darmstadt, Germany), anti-APLP2 (Cat. No. 171616, Calbiochem EMD Biosciences – Merck, Darmstadt, Germany), anti-GAPDH (14C10, Cell Signaling, Danvers, MA, USA), anti-MAP2 (M4403, Sigma-Aldrich St Louis, MO), anti-GAD65 (D5G2, Cell Signaling), anti-mouse IgG, HRP Whole antibody (NA931-1ML, Amersham, Little Chalfont, UK), anti-rabbit IgG, HRP Whole antibody (NA934-1ML, Amersham, Little Chalfont, UK), goat anti-mouse Alexa Fluor®-488, goat anti-mouse Alexa Fluor®-568, goat anti-rabbit Alexa Fluor®-647 and DAPI were purchased from ThermoFisher Scientific (Waltham, MA, USA). Glutamate assay kit was from Abcam (Cambridge, UK) and γ-aminobutyric acid (GABA) ELISA was purchased from Cloud-Clone Corporation. 70µm Falcon™ Cell Stainers were from ThermoFisher Scientific (Waltham, MA).

### Animal models

APP+/+ and APP-/- mice were obtained from the Jackson Laboratory (Bar, Harbor, ME, USA) as C57Bl6/J and backcrossed for > 6 generations in CD1 genetic background. Animals were housed on a 12 h light/dark cycle in standard animal care facility with access to food and water *ad libidum*. Heterozygous animals (APP+/-) were bred and crossed to obtain embryos from the three different genotypes (APP+/+, APP+/- and APP-/-) in the same litter. All experiments were performed in compliance with protocols approved by the UClouvain Ethical Committee for Animal Welfare (code number 2016/UCL/MD/015).

### Primary neurons culture and treatments

Primary cultures of cortical neurons were prepared from E18 mouse embryos as previously described (Pierrot et al., 2013). Briefly, cortices were dissected and dissociated in HBSS without calcium and magnesium and the mixture was centrifuged on Fetal Bovine Serum (FBS) for 10 min at 1000xg to pellet cells. Cells were plated at 200.000 cells/cm^2^ in culture dishes pre-treated with 10 µg/ml of poly-L-lysine in phosphate buffered saline (PBS) and cultured for 3 to 14 days *in vitro* in Neurobasal® medium enriched with 2% v/v B-27® supplement medium and 1mM L-glutamine at 37°C, 5% CO_2_ and humidified atmosphere. Half of the medium was renewed every 2-3 days.

After 6 days (DIV6), neurons were treated for 16h with 1µM of N-[N-(3,5-Difluorophenacetyl)-L-alanyl]-S-phenylglycine t-butyl ester (DAPT), a γ-secretase inhibitor (Dovey et al., 2001) or with 20 nM of soluble APP alpha in Neurobasal® Medium

### Primary astrocytes culture and treatments

Cortices from rat pups were collected at postnatal day 2 and mechanically dissociated. Astrocytes were isolated using a 30% Percoll gradient and seeded into gelatin-coated tissue culture flasks. ^Cells were left to proliferate for 14 days at 37°C - 5% CO^2 ^in DMEM-glutaMAX medium^ supplemented with 10% FBS, 50 mg/ml penicillin–streptomycin and 50 mg/ml fungizone. Medium was renewed after 7 days, cells were passaged after 14 days and further cultured in DMEM-glutaMAX with 10% FBS. Two days later, FBS was reduced to 3% and medium was supplemented with the growth factor cocktail G5. All experiments were conducted 7 days later (DIV7).

### Npas4 induction analysis

For Npas4 induction analysis, neurons and astrocytes at DIV7 were depolarized with 50mM potassium chloride. Cell lysates were analyzed by Western blotting with the anti-Npas4 antibody. as described below.

### RNA extraction, transcriptome analysis and qRT-PCR

Total RNA was extracted by TriPure Isolation Reagent according to the manufacturer’s protocol. RNA samples were suspended in DEPC-treated water and RNA concentration was measured (OD 260 nm) on BioSpec-nano spectrophotometer (Shimadzu Biotech). For microarray analysis, RNA quality was evaluated by capillary electrophoresis using the Agilent 2100 Bioanalyzer instrument with the Agilent RNA 6000 Nano Kit according to the manufacturer’s instructions (Agilent, Santa Clara, CA). 250 ng of total RNA for each sample was amplified and labeled using GeneChip®WT PLUS Reagent kit (Affymetrix) before being hybridized on *GeneChip®Mouse Transcriptome 1.0 Array*, overnight at 45°C. The chip was washed using an automated protocol on the GeneChip® Fluidics Station 450 followed by scanning on a GeneChip® Scanner on Affymetrix microarray platform (de Duve Institute, UCL, Brussels).

For quantitative PCR, RNA samples were reversed transcribed using iScript cDNA Synthesis Kit and real time PCR was performed in an iCycler MyIQ2 multicolor-Real-Time PCR detection system using iQ SYBR Green supermix kit (Biorad). A standard curve was established for relative quantification with a fourfold dilution series (from 100 to 0,0097 ng) of a cDNA template mix. Relative quantification was calculated by the 2^ΔΔCT^ method, and results were normalized first to *Gapdh* expression and then normalized (percentage or fold) to the control condition. Primers used are depicted in Table 1 in Supplementary material.

### Western blotting

Cells were solubilized and sonicated in lysis buffer (20% Glycerol, 4% SDS, 125 mM Tris-HCl pH 6.8) containing a cocktail of proteases and phosphatases inhibitors. Mice were euthanized (Ketamine/Xylazine injection) and brains were dissected after perfusion with ice cold sterile PBS. Cortices and hippocampi were isolated and quickly frozen in liquid nitrogen until use. Tissues were crushed using mortar pestle method. For brain protein extraction, samples were homogenized in RIPA buffer (1% (w/v) NP40, 0.5% (w/v) deoxycholic acid, 0.1% (w/v) SDS, 150 mM NaCl, 1 mM EDTA, 50 mM Tris, pH 7.4) containing proteases and phosphatases inhibitors cocktail (Roche, Basel, Switzerland). The samples were clarified by centrifugation at 20,000 x g. Protein concentrations were determined with a BCA kit. Samples were prepared with NuPAGE LDS sample buffer (4x) and 50 mM DTT and then heated for 10 min at 70°C. 10 to 40 µg of proteins or 22 µl of culture medium were loaded per well for migration followed by transfer onto PVDF or nitrocellulose membranes. For APP C-terminal fragments blotting, proteins were transferred on nitrocellulose (0.1 µm). Membranes were blocked in nonfat dry milk (5% in PBS, 0,1% Tween-20) and immunoblotted with anti-APP NT (22C11, 1/500), anti-APP CT (Y188, 1/500), anti-APLP1 (1/1000), anti-APLP2 (1/1000) and anti-GAPDH (1/25000). Blots were revealed using ECL and signal quantification was performed using GelQuant.NET software (BiochemLabSolutions.com).

### ImmunoCytoFluorescence (ICF)

Neurons were grown at 100.000 cells/cm^2^ per well on poly-L-lysine coated coverslips. Neurons were rinsed with PBS and fixed for 15 min in PBS/4% paraformaldehyde. Neurons were washed again twice in PBS for 5 min and processed as described previously (Decock et al., 2016). Permeabilization and blocking steps were done in PBS/5% skimmed milk/0.3% Triton-X100 (M3TPBS); antibodies were incubated in PBS/5% skimmed milk/0.1% Triton-X100 (M1TPBS). Primary antibodies dilutions used: mouse anti-MAP2 (1/1000), rabbit anti-APP (Y188, 1/100) and rabbit anti-GAD65 (D5G2, 1/100). Secondary antibodies dilutions used: goat anti-mouse Alexa Fluor®-488 (1/500), goat anti-mouse Alexa Fluor®-568 (1/500) and goat anti-rabbit Alexa Fluor®-647 (1/500). Images were acquired on Evos FL Auto microscope (Invitrogen) with GFP (Alexa Fluor®-488 or native GFP), TxRed (Alexa Fluor®-568) and CY5 (Alexa Fluor®-647) EVOS LED light cubes and analyzed with ImageJ software. For the quantification of signal area, 10X or 20X magnification images were identically thresholded for APP+/+ and APP-/- or Ct and CRISPR-*Npas4*. Area of thresholded images was measured and normalized to the number of cells counted by DAPI staining. For the quantification of the APP expression intensity, image acquisition was performed using 40x objective coverslip-corrected (ThermoFischer Scientific, AMEP4699) in GFP, CY5 (APP) and DAPI channels. A total of 12, 19 and 19 images were acquired to obtained 33, 46 and 51 neurons in the analysis (Figure 3B) for CRISPR control (Ct), Oligo2 and Oligo17 respectively. GFP channel images were first 8-bit transformed and thresholded to highlight only GFP staining. A region of interest (ROI) was delimited around GFP+ neurons in the GFP channel (green using “wand tool” in imageJ software and transposed to CY5 (APP) channel (blue). ROI mean intensity is measured using “Analyze” tool of ImageJ software.

### AICD and CRISPR/Cas9 lentiviral constructions and production

We used a lentiviral vector-based approach to express AICD in neurons. AICD50 tagged at the c-terminal part with hemagglutinin (HA) was cloned into pLenti CMV/TO Puro lentiviral vector (Addgene #17482). pLenti CMV/TO Puro empty is used as control (Ct). We used a lentiviral vector-based approach to deliver the CRISPR-Cas9 system. We designed sgRNAs “Oligo2” and “Oligo17” to target *App* mouse gene (Gene ID: 11820), and sgRNA “CRISPR-*Npas4*” to target Npas4 mouse gene (Gene ID: 225872). sgRNAs were cloned in a lentiviral vector delivering sgRNA, SpCas9 and coexpressing eGFP (Addgene #57818) according to author instructions (Heckl et al., 2014). The negative control (Ct) used was the lentiviral construct without sgRNA but expressing SpCas9 and eGFP. sgRNA sequences, scores and PAMs are provided in Table 2 in Supplementary material. Briefly, sgRNAs purchased at Sigma-Aldrich (St Louis, MO, USA) were designed using on/off-target score algorithm and cloned into the pL.CRISPR.EFS.GFP plasmid. Vectors were validated by sequencing (Beckman Coulter Genomics, UK), produced and purified using Plasmid Midi kit (Qiagen, Hilden, Germany). Lentiviruses were produced by transfecting HEK293-T cells in 10 cm dishes (2×10^6^ cells/dish) with lentiviral CRISPR-Cas9 vectors, pCMV-dR8.2 (Addgene#12263) and pMD2.G (Addgene#12259). After 48 h, the supernatant was filtered and incubated with 1/3 (v/v) of LentiX™ Concentrator for 90 min on ice. The collected supernatant was centrifuged at 1500xg for 45 min at 4°C, the pellet was resuspended in 20 µl per dish of Neurobasal® Medium and stored at -80°C until use. Empty backbone of pL-CRISPR.EFS.GFP was used as negative control (Ct) in our studies.

Neurons were infected with lentiviruses CRISPR-Cas9 1 day after plating (DIV1). Typically, 20 µl of concentrated virus were used to infect 800.000 cells per well of 12 well plate dish. The medium was completely changed after 24 hours and a half media change was carried out every 2-3 days thereafter. The neurons were harvested at 7 days in vitro (DIV7) or as indicated.

### Lentiviral toxicity assay

Cell viability was measured by LDH release in the culture medium at DIV7 after lentiviral infection using Cytotoxicity Detection kit (Sigma-Aldrich, St-Louis, MO, USA) according to the manufacturer's instructions. Relative absorbance was measured at 490 nm using a VICTOR Multilabel Plate Reader (PerkinElmer, Richmond, VA, USA). Background LDH release was determined in non-infected control cultures.

### Flow cytometry and cell sorting

After DIV7, infected neurons were briefly rinsed with PBS and trypsinized for 2 min. Neurons were mechanically dissociated and filtered through 70 µm Falcon™ Cell Strainers in 50 ml tube containing FBS. Cells were pelleted by centrifugation at 1000xg for 5 min and resuspended in PBS/1% FBS/1mM EDTA. TO-PRO™-3 Iodide (Thermo Fisher Scientific) was used to stain dead cell and exclude them for the sorting. Cells were sorted using a BD FACSAria™III cell sorter (BD Biosciences, San Jose, CA) on the “Flow cytometry and cell sorting - CYTF” UCL platform. The sort parameters used were the following: nozzle 100 µm, sheath pressure 20 psi, drop frequency 30 kHz and sort precision 16-32-0. Sample and collection tubes were maintained at 4°C throughout the sort. GFP-negative and positive cells were harvested in PBS/1% FBS/1mM EDTA and centrifuged at 12000xg for 2 min and homogenized in TriPure Isolation Reagent for RNA extraction.

### Glutamate and GABA measurements

Glutamate and γ-aminobutyric acid (GABA) were measured in medium and in cells at DIV7. Briefly, neurons were grown at 200.000 cells/cm^2^ in 12 well plate culture dish. Media were harvested, centrifuged to pellet cell’s debris and supplemented with cocktail of proteases inhibitors and frozen at -20°C until use. Cells were scratched in ice cold PBS and pelleted by centrifugation (12.000xg for 3 min at 4°C) then quickly frozen in liquid nitrogen and kept at -80°C until use. *For glutamate assay*: Media were directly used as are. Cells were prepared according to the manufacturer protocol and measurement was normalized on protein content. *For GABA ELISA assay:* Media were directly used as are. Cells were lysed by 5 cycles of thawing and freezing in PBS and centrifuged at 12.000xg for 10 min at 4°C. Supernatant was used for the quantification and normalized on protein content.

### Statistical analysis

*Microarray analysis*: Raw data were analyzed using Bioconductor (R environment). Robust Multiarray Average (RMA) was used for background correction, normalization, probe level intensity calculation and probe set summarization. Gene expression values were compared between APP+/+ and APP-/- neurons at different stage of development DIV3, DIV7 and E18 using the R-Limma (Linear Models for MicroArray Data) package. Benjamini-Hochberg procedure was used for multiple testing corrections. From raw data, only transcripts with an Entrez ID were kept in order to facilitate the analysis. Gene set enrichment analysis was performed on differentially expressed genes sets after the ROAST (Rotation gene set tests for complex microarray experiments) (Wu et al., 2010) procedure to identify KEGG pathways modified in absence of APP for all conditions (E18, DIV3 and DIV7). The data obtained have been deposited in NCBI's Gene Expression Omnibus (Edgar et al., 2002) and are accessible through GEO Series accession number GSE112847 (https://www.ncbi.nlm.nih.gov/geo/query/acc.cgi?acc=GSE112847). Otherwise, statistical analyses were performed using GraphPad Prism (GraphPad Software, San Diego, CA). Gaussian distribution was assessed by Kolmogorov-Smirnov test (GraphPad Prism). If the data follow normal distribution parametric test was applied. Otherwise non parametric test was used. If two groups were compared, parametric Student’s t-test or non-parametric Mann-Whitney test were used. If more than two groups were compared, parametric ANOVA with post hoc tests as indicated or non-parametric Kruskall-Wallis were used. (*, p < 0,05; **, p < 0,01; ***, p < 0,001). The number of biological replicate (n) analyzed is indicated in figure legends in the number of independent experiment (N).

## Results

### APP-dependent expression of *Npas4* in differentiated primary neuron cultures

Experiments were performed on primary neuron cultures according to the workflow described in Supplementary Figure S1A. Briefly, neurons from embryonic cortex (E18) were cultured for 3 or 7 days in vitro (DIV3 or DIV7) and longer (up to DIV14) when necessary. We first characterized the expression of APP family proteins and observed an increase in APP, APLP1 and APLP2 upon differentiation with a peak of expression at DIV 7-8 (Figure S1B-C), supporting an important role of APP protein family in neuronal maturation. No modifications of APLP1 or APLP2 levels and maturation were observed in APP-/- neurons when compared to APP+/+ at any time point of differentiation studied (Figure S1D). Thus, results obtained here in APP-/- neurons can be related to the loss of APP and not to indirect effects resulting from up- or down-regulation of APLP1 or APLP2. Previous studies indicated that AICD is detectable inside the nucleus specifically at DIV6-7 (Kimberly et al., 2005) suggesting that AICD-dependent gene transcription is temporally restricted. We checked for AICD production at DIV7 in total lysates of APP+/+, APP+/- and APP-/- cultures (Figure S1E). AICD was readily detectable in APP+/+ neurons at but only at high exposure time, confirming that it is a transient peptide (Huysseune et al., 2007) with a restricted temporally expression pattern in primary neurons. AICD-dependent transcriptional regulation may therefore only occur within a defined time-period, around DIV7 (and not at DIV3).

To investigate this, we performed microarray experiments, which allow description of genome-wide expression changes in APP +/+ and APP-/- primary cortical neurons at DIV3 (immature neuronal network), DIV7 (neuronal network with detectable AICD) and in E18 cortical tissue (summarized in Figure S1A). We used *Affymetrix GeneChip®Mouse Transcriptome 1.0 Array* and carried out data analysis with the R-Limma (**Li**near **M**odels for **M**icro**A**rray Data) package (Ritchie et al., 2015). The chips used allow the profiling of coding and non-coding (lncRNA, miRNA, pseudogene…) gene expression as well as alternative splicing events. We ran each condition (E18, DIV3, and DIV7) in triplicate (3 chips used for each condition, from independent cultures). We focused here on differentially expressed coding genes, although data were collected for non-coding RNAs (not shown). Strikingly, the overall changes observed (fold changes) were moderate in all conditions (E18, DIV3 and DIV7). Few coding transcripts appear to be differentially expressed when the specific fold change (linear) is set at 1.25, 1.5 or 2 (Figure 1A). The Benjamini-Hochberg multiple correction test did not reveal any robust differential gene expression (adjusted p-value <0,05) except as expected for *APP* (positive control). Gene enrichment analysis was performed using ROAST (Rotation gene set test for complex microarray experiment) procedure to finally identify molecular interaction/reaction networks diagram (Kanehisa and Goto, 2000) also known as KEGG pathway altered in the absence of APP. The first five pathways (in terms of significance), the number of genes modified as well as their direction are shown in Figure 1B. Interestingly, *ECM (extracellular matrix)-receptor interaction* and *Long-term potentiation* pathways are modulated in absence of APP at DIV7. Cell-ECM interactions are mediated by transmembrane receptors and cell adhesion proteins, involved in adhesion, differentiation and maturation. Long term potentiation (LTP) is a major mechanism in memory formation and learning. Both of these pathways have been associated to APP function (Caceres and Brandan, 1997; Seabrook et al., 1999; Puzzo et al., 2011). To note, we did not measure any expression change (Supplementary Table 3) of genes identified as AICD target genes (Pardossi-Piquard and Checler, 2012).

**Figure 1:**
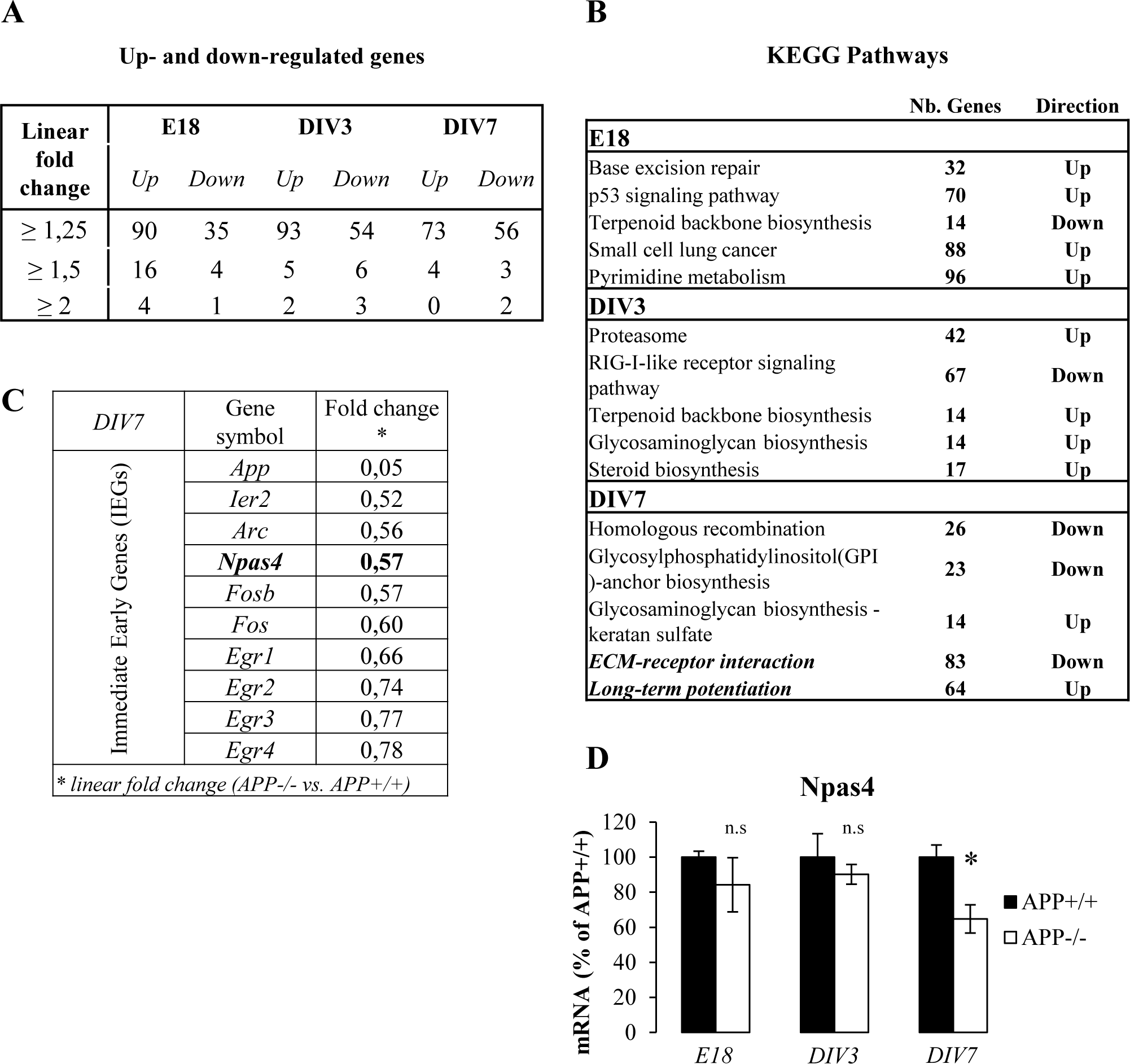
**APP-dependent expression of *Npas4* in young differentiating neuronal culture Summary of transcriptome analysis performed with the *GeneChip® Mouse Transcriptome Array 1.0* (Affymetrix).** Data were processed in triplicate (3 independent cultures) for each experimental time point (E18; DIV3; DIV7). Non-coding transcripts and alternative splicing products are detected by these arrays, but only transcripts of coding transcripts have been considered here. **A**) Number of up-and down-regulated coding transcript in APP-/- vs. APP+/+ primary neurons at E18, DIV3 and DIV7. Linear fold changes have been set at 1,25, 1,5 and 2. **B**) KEGG pathway analysis (http://www.genome.jp/kegg/pathway.html) at E18, DIV3, DIV7 (APP-/- vs. APP+/+) to identify networks molecular pathways (or interaction networks) in which differentially expressed genes are clustered. The five most modified pathways are displayed for each time point, with the number of genes potentially up-or down-regulated. **C**) Immediate Early Genes (IEGs) expression in APP-/- vs. APP+/+ primary neurons at DIV7 and their respective fold change (APP-/- vs APP+/+) in microarray analysis at DIV7. **D**) Neuronal PAS 4 domain (Npas4) mRNA level was measured by qPCR at E18, DIV3 and DIV7 (n=6, N=3). Results (mean ± SEM) are expressed as percentage of controls (APP+/+). n.s.= non-significant, *p=0.0242, Student’s t-test.

We decided to further select genes relevant to described APP cellular function in order to further investigate their regulation by APP. In a set of array (APP+/+, accession number GSM3089741 vs APP-/- accession number GSM3089744) from a primary neurons at DIV7, we noticed a down-regulation of (IEGs) in APP-/- neurons (Figure 1C). Among them, the activity-dependent transcription factor, *Npas4* (Neuronal PAS domain protein 4) was of particular interest. NPAS4 is a neuron-specific IEG, known to be regulated by neuronal activity and involved in synaptic plasticity and synaptic homeostasis. We confirmed by qPCR that the *Npas4* mRNA level was decreased at DIV7 in APP-/- neurons compare to APP+/+, but not at DIV3 nor in the cortex at E18 (Figure 1D). To note, the expression of other early genes (Egr1 and Egr3) previously reported to be involved in APP-dependent gene expression (Hendrickx et al., 2013) were not altered in our conditions (Figure S2A, S2B).

### *Npas4* expression is AICD-independent

Since transcription of APP target genes could involve AICD (Belyaev et al., 2010), which is produced particularly at DIV7, we inhibited its release by treating neurons with 1µM DAPT for 16 h, a well-described non-competitive γ-secretase inhibitor (Dovey et al., 2001). DAPT treatment induced APP CTFs accumulation (Figure 2A), indicating that γ-processing thus AICD release were inhibited under these conditions (Hage et al., 2014). DAPT treatment didn’t decrease Npas4 expression in APP-/- primary neurons, but indeed increased it in APP+/+. To further address the role of AICD in Npas4 regulation, we transduced primary neurons with a lentiviral vector expressing the 50 C-terminal amino acids of APP (AICD) fused (C-terminus) to the hemagglutinin tag (HA). AICD-HA is detectable in infected cells (Figure 2C), and AICD expression in APP-/- neurons did not modify *Npas4* mRNA levels (Figure 2D), confirming that AICD is not involved in APP-dependent *Npas4* transcriptional regulation. As some of the APP functions were found to rely on its extracellular soluble fragment (sAPPα), we tested whether the sAPPα can regulate *Npas4* expression *per se*. Neuronal cultures were treated with 20 nM of human sAPPα for 16 h (Figure 2E) and *Npas4* expression was measured by qPCR. *Npas4* mRNA levels increased significantly upon sAPPα addition in APP+/+ neurons, but not in APP-/- (Figure 2F). These data provide a general insight into APP-dependent *Npas4* transcription in neurons. (i) AICD release is not involved in this process; (ii) APP soluble ectodomain (sAPPα) regulates *Npas4* expression, only in a context where endogenous APP is expressed. Important to note, glial cells represent about ~16% of total cells in primary cultures, and can indirectly contribute to the mechanisms we observed. However, absence of APP did not change the astrocytic pattern of primary cultures, and astrocytes do note express readily detectable Npas4 levels (Figure S3A-B). Hence, our observations reflect APP-dependent Npas4 regulation truly acting in neurons

**Figure 2:**
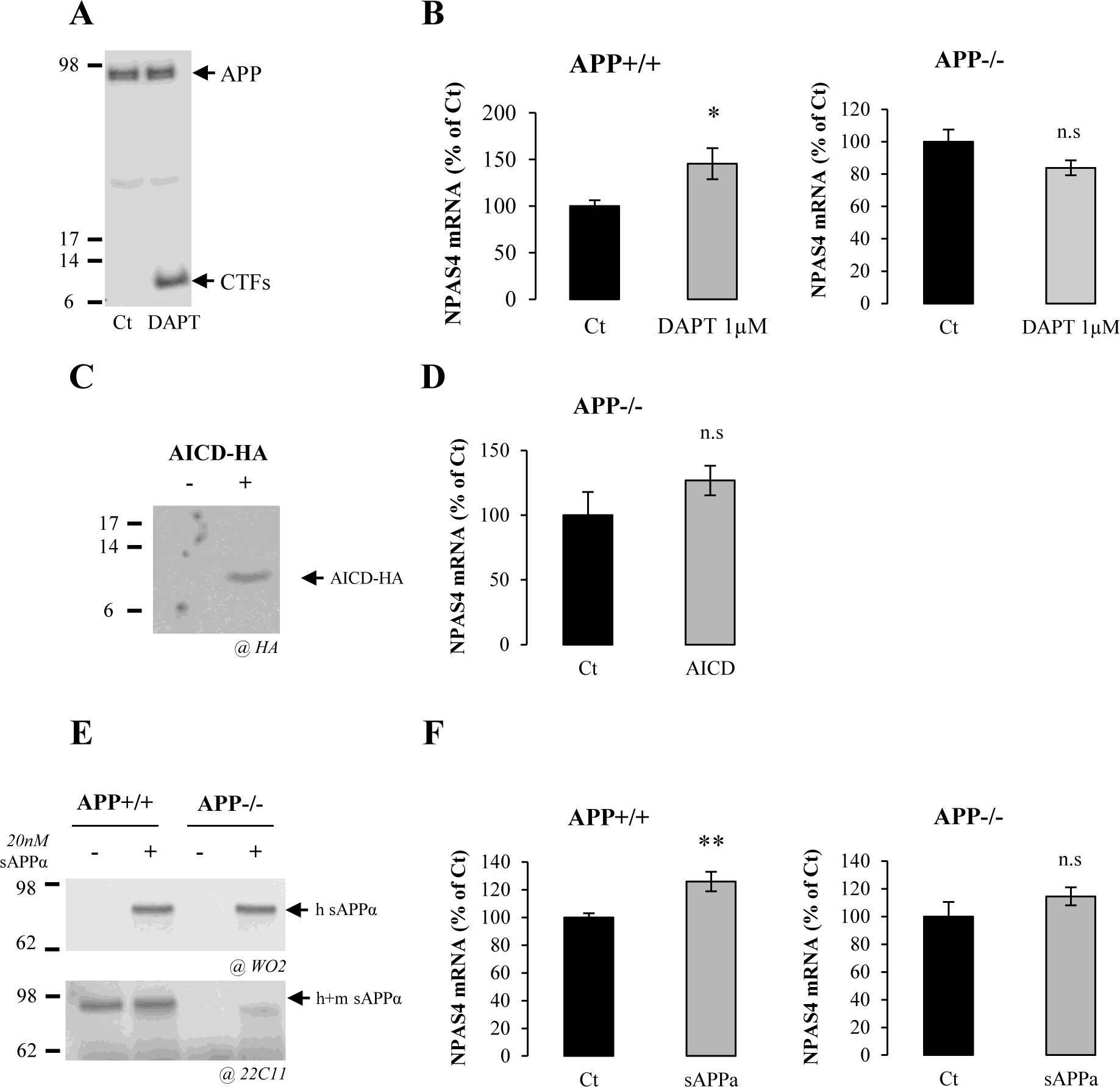
**APP metabolites but not AICD regulate *Npas4* expression A) Western blotting analysis of APP and C-terminal fragments (CTFs and AICD) in cortical neurons after 1 µM DAPT treatment for 16 h at DIV7. B**) Quantification by qPCR of *Npas4* mRNA in APP+/+ or in APP-/- neurons at DIV7 treated with 1 µM DAPT for 16 h (n=7, N=4). Results are expressed as percentage of control (Ct) (mean ± s.e.m.). *p=0.0216, n.s.= non-significant, Student-t test. **C**) Western blotting analysis of AICD-HA expression after 3 days of lentiviral infection in cells with control or AICD-HA expressing vectors. Total cell lysate was analyzed with anti-HA antibody. **D**) Quantification by qPCR of *Npas4* mRNA in neurons at DIV7 infected with lentiviral vector expressing AICD-HA (n=6, N=2). Results are expressed as percentage of control (Ct) (mean± s.e.m). n.s.= non-significant, Student-t test. **E**) Medium of sAPPα treated APP+/+ or APP-/- neurons was subjected to western blotting analysis using anti-human APP antibody (clone WO2) to detect the exogenous human sAPPα (h sAPPα) and using anti-mouse APP antibody (clone 22C11) to detect both endogenous and exogenous sAPPα (h+m sAPPα). Medium was collected after 16 h of treatment. **F**) Quantification by qPCR of *Npas4* mRNA level in APP+/+ (n=8, N=4) or in APP-/- neurons at DIV7 treated with 20 nM sAPPα for 16 h (n=6, N=3). Results are expressed as percentage of control (Ct) (mean ± s.e.m.). **p=0.0055, n.s.= non-significant, Student-t test.

The APP-dependent transcriptional regulations we observed were subtle when compared to those reported in the literature, in line with the mild phenotype of APP knockout mice (Muller et al., 1994; Zheng et al., 1995). Although we did not observe compensation of APP loss by APLP overexpression in our model, APP-dependent gene regulations that appear in the close-up could be hidden in the long term or related to functional redundancies with other members of the APP family (Shariati and De Strooper, 2013). In this line, APP-/- mice brain phenotype is better unraveled by acute down-regulation of APP (Senechal et al., 2007). We decided therefore to knock-down the APP expression in APP+/+ neurons with a lentiviral-based CRISPR-Cas9 genome editing approach (Jinek et al., 2012), to test the consequence of acute APP knock-down on *Npas4* expression. Nearly ~50% of the cells in culture were infected under our conditions (Figure S4A-B) and no lentiviral toxicity was measured (Figure S4C). Only neuronal cells were infected, reflecting the tropism of the viral particles for neurons, and not for glial cells. APP expression was monitored by ICF (Figure 3A) and by measuring the intensity of APP signal in GFP-positive (infected) neurons (Figure 3B). APP was strongly decreased in neurons infected with viruses expressing the Oligo2 and 17 sgRNA sequences targeting APP exon1 and exon 2, respectively, when compared to Ct. This was confirmed by Western blotting showing the APP expression specifically decreased by about 50%. Importantly, APLP1 and 2 expressions were not altered in cultures infected with Oligo2 and Oligo17 lentiviruses (Figure 3 C), clearly indicating that off targets mechanisms - a major risk with CRISPR-Cas9 approaches, especially with homologous genes - are not observed in our experimental setup.

**Figure 3:**
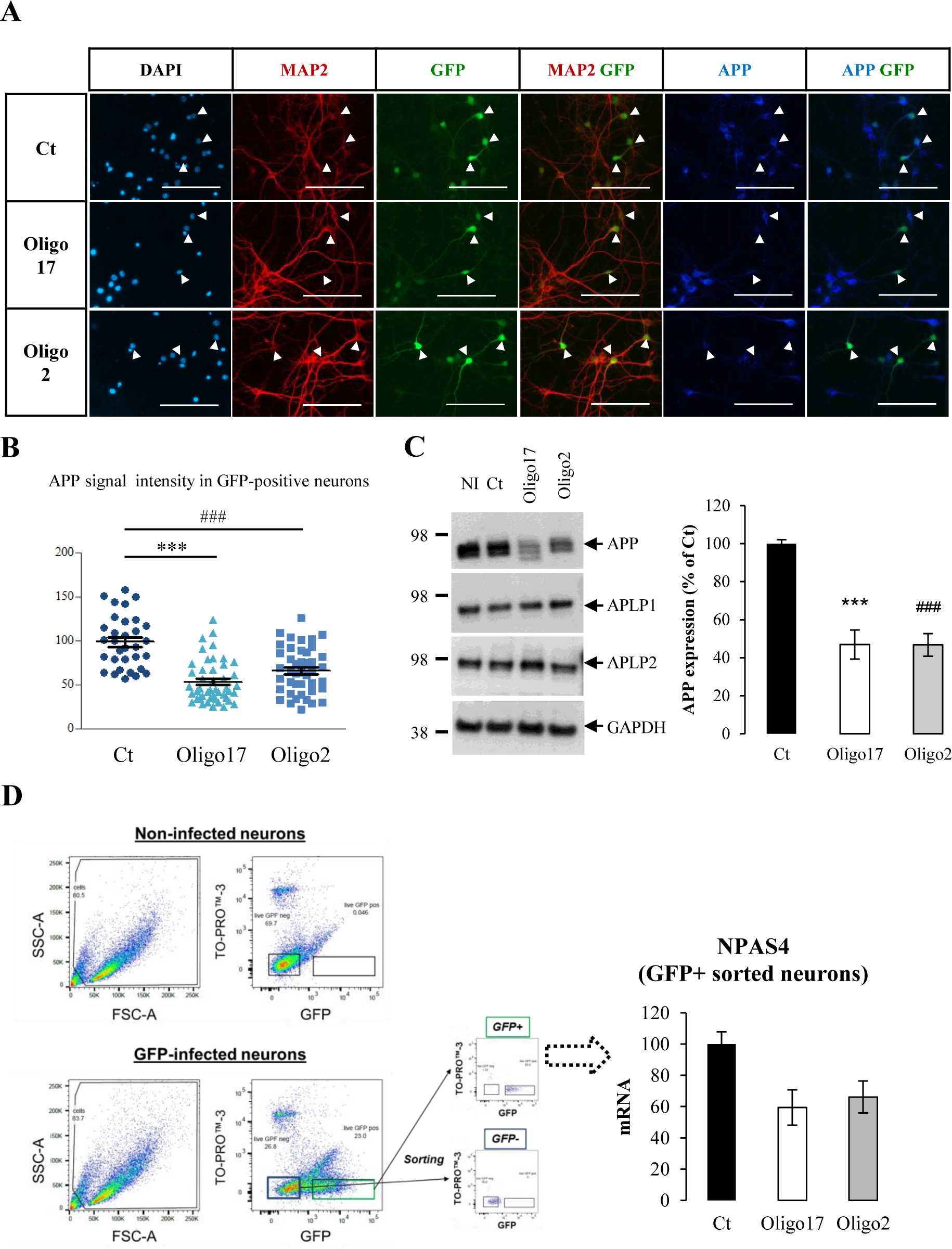
**Decreased *Npas4* expression in APP-silenced primary neurons APP was knock-down by CRISPR-Cas9 approach in primary neurons cultures. A**) Cortical neurons were infected at DIV1 with lentiviruses expressing sgRNAs (Oligo2, Oligo17) or no sgRNA (Ct), SpCas9 and GFP. Cultures were immunostained for MAP2 (red), APP (blue) and DAPI (light blue) at DIV7. Arrowheads indicate the position of GFP-positive (infected) neurons in each condition. Scale bar: 100µm. **B**) Quantification of APP signal in GFP-positive neurons. At least 33 neurons were quantified in two independent experiments for each condition (n=33 N=2). Results (mean ± SEM) are given as percentage of control (Ct). ###p<0,001 (Ct vs Oligo 2) ***p<0,001 (Ct vs Oligo17); Kruskal-Wallis test and Dunn’s multiple comparison test. **C**) Left panel. Representative Western blots showing APP, APLP1, APLP2 and GAPDH protein level in cortical neurons at DIV7 infected in the same conditions. NI = non-infected. Right panel, quantification of APP expression measured by Western blotting. Results (mean ± SEM) are given as percentage of control (Ct). ***p<0,001 (Ct vs Oligo17), ###p<0,001 (Ct vs Oligo 2), ANOVA and Bonferroni’s multiple comparison test (n=6, N=3). **D**) Sorting of GFP-expressing neurons (FACS). Scatter plots (FSC vs. SSC, left panels) of non-infected and GFP-expressing cells are shown. Dot plots (TOPRO-3, far red vs. GFP, right panels) were used to gate (green rectangle) GFP-positive/TOPRO-3 negative cells. RNA was extracted from these cells and Npas4 mRNA level was quantified by qPCR. Results were obtained from pooled samples (4 wells of 4 cm^2^ each) for each condition (Ct, Oligo2 and Oligo17). Quantification were carried out on 2 independent experiments (N=2). Results (mean ± SEM) are expressed as percentage of Ct.

We decided to measure the expression of *Npas4* after APP knock-down selectively in GFP-positive (knock-down) neurons. This was achieved by sorting GFP positive cells by flow cytometry. We used TO-PRO™-3 staining as a viability marker to exclude dead cells from the analysis and set sorting parameters by using non-infected condition (no GFP) and neurons expressing GFP (GFP infected) as standards. *Npas4* mRNA levels were measured in these cells by qPCR (Figure 3D). *Npas4* mRNA was readily decreased in neurons infected with Oligo2- and 17-expressing lentiviruses. Acute APP knock-down achieved with the CRISPR-Cas9 system in primary neurons resulted in the decrease in *Npas4* expression, confirming the APP-dependent *Npas4* transcriptional expression observed in APP deficient neurons.

### APP deficiency increases the markers of GABAergic transmission

Down-regulation of *Npas4* expression in the absence of APP could reflect an impairment in neurite formation and/or synaptogenesis which may lead to deficient in basal neuronal activity. APP was reported to modulate neurite outgrowth and synapse formation (Priller et al., 2006; Young-Pearse et al., 2007; Tyan et al., 2012; Billnitzer et al., 2013) but the mechanisms by which APP modulates synapse formation and plasticity is poorly understood. We first analyzed neuronal arborization at DIV7, when Npas4 expression is decreased. We monitored arborization by measuring the area of the neuron-specific microtubule associated protein2 (MAP2) signal per cell from DIV1 to DIV7 (Figure 4A). APP-/- neurons extend neurites and no difference was observed at DIV1 to DIV3. The absence of APP subtly (but significantly) increased MAP2 signal at DIV7 (Figure 4B), indicating the importance of APP for proper neurite arborization.

**Figure 4:**
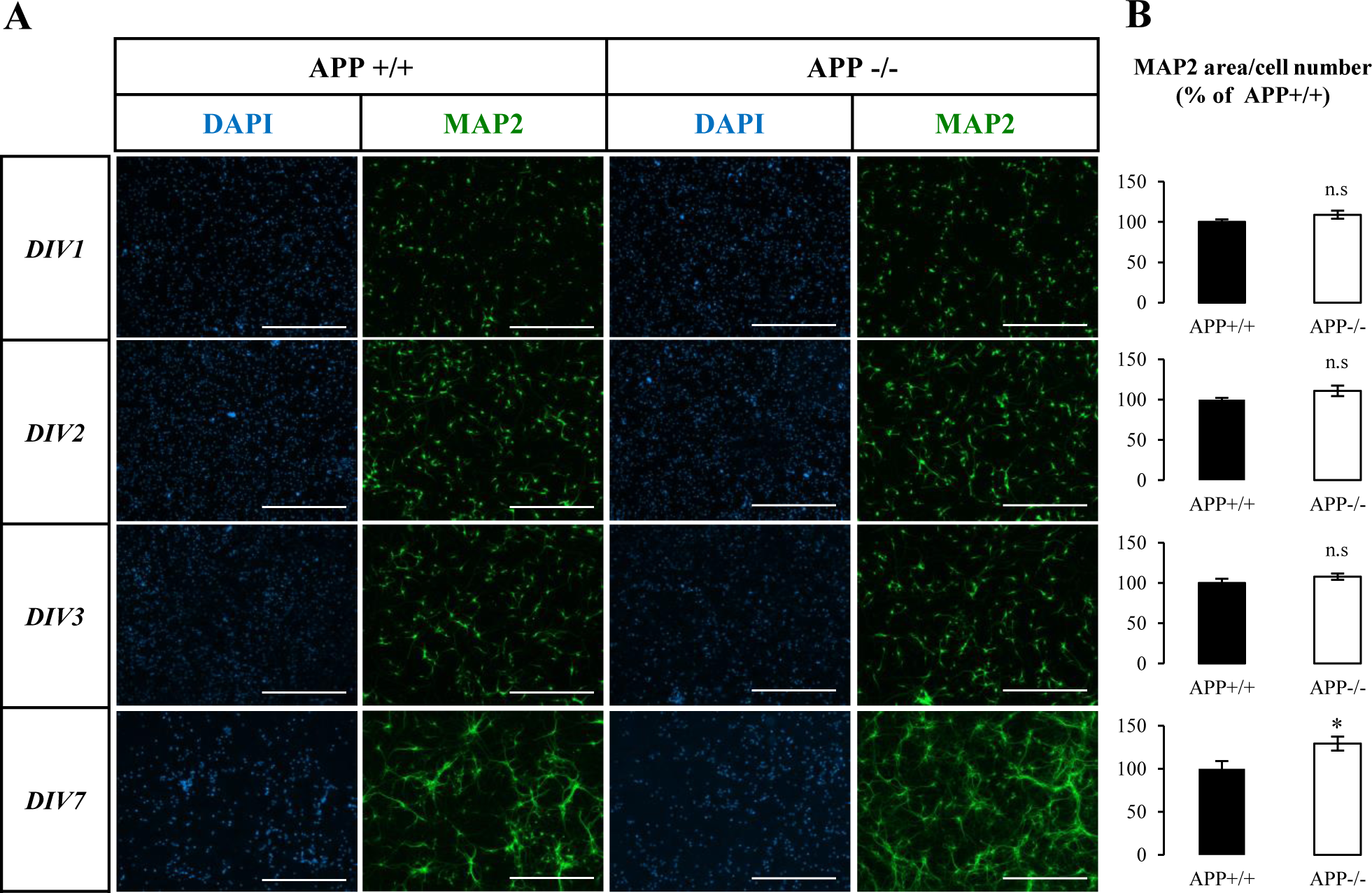
**Altered neurites arborization of APP deficient neurons during in vitro maturation A**) Cortical APP+/+ or APP-/- were stained against the neuron-specific marker MAP2 and the nuclear dye DAPI at different stages of maturation (DIV1-2-3 and DIV7). Scale bar: 400µm. **B**) Quantification of MAP2 signal area normalized to the number of neurons at DIV1, 2, 3 and 7. Quantifications were from 3 fields of at least 6 coverslips from APP+/+ and APP-/- neurons, in three independent experiments (N=3). Results (mean ± SEM) are expressed as percentage of control (APP+/+). *p=0,0293, Mann-Whitney test.

*Npas4* is involved in the fine tuning of excitatory/inhibitory homeostasis, by controlling the balance of excitatory and inhibitory inputs on post-synaptic neurons (Lin et al., 2008; Bloodgood et al., 2013; Spiegel et al., 2014). This characterized by the type of neurotransmitter released: typically glutamate for excitatory synapses and GABA for inhibitory synapses. We measured the amount of GABA and glutamate released in medium and present in the cells at DIV7 (Figure 5A-B). The concentration of GABA is increased by 83% in the medium of APP-/- neurons (Figure 5A), and no difference was observed for GABA measured in cells. Strikingly, we observed no significant change in glutamate concentration (cell or medium) in APP-/- neuronal cultures compare to APP+/+ (Figure 5B). This is supported by the only slight qualitative modifications in glutamate responses measured by intracellular calcium imaging in APP-/- neurons (Figure S5). GABA is ^synthetized by the glutamate decarboxylase enzymes (GAD^65 ^and GAD^67^) that catalyze the decarboxylation of glutamate to GABA. GAD^65 ^synthesizes GABA for neurotransmission, and is therefore active at nerve terminals and synapses. By immunostaining, we observed that GAD^65 signal is increased in APP-/- neurons compared to APP+/+ neurons (Figure 5C). This is not caused by an increase in the relative number of GABAergic neurons in APP-/- cultures compare to APP+/+ (Figure S6), pointing to an increase in GAD65 cellular expression. To further address the effect of APP deficiency on GABAergic neurotransmission, we quantified the expression of the most prevalent GABA receptor subunit, GABARa1 expressed during neuronal development. We found GABARa1 to be slightly but significantly decreased in APP-/- neurons (Figure 5D), suggesting complex modifications of GABAergic neurotransmission. In summary, our results indicate that APP deficiency disturbs mainly GABAergic neurotransmission components with a little effect on the excitatory counterpart.

**Figure 5:**
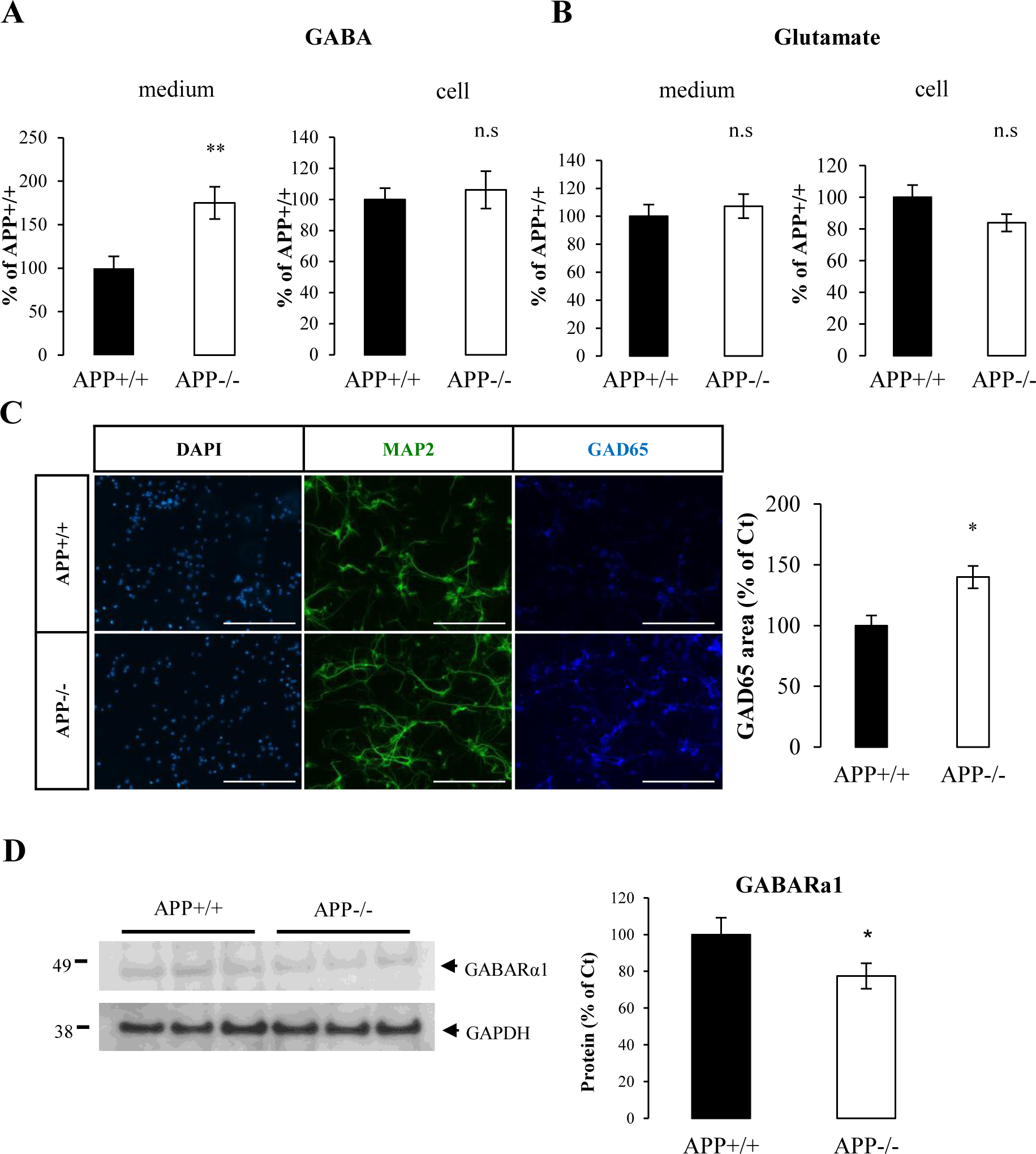
**GABAergic markers are impaired in APP knock-out neurons A**) Quantification of γ-amino butyric acid (GABA) in culture medium and cell extracts of APP+/+ and APP-/- primary neurons at DIV7. Results (mean ± SEM) are expressed as percentage of APP+/+ (n=20, N=3). **p=0,0024, n.s.= non-significant, Student-t test. **B**) Quantification of glutamate in culture medium and cell extracts of APP+/+ and APP-/- neurons at DIV7. Results (mean ± SEM) are expressed as percentage of APP+/+ (n=16, N=3). n.s.= non-significant, Student- t test. **C**) Cortical APP+/+ and APP-/- neurons at DIV7 were immunostained against the neuron-specific marker MAP2 and glutamate decarboxylase 65 (GAD65). Quantification of GAD65 signal area (5 fields per coverslip) was normalized to the number of cells. At least 2 coverslips were quantified for each group (APP+/+ and APP-/-) in two independent experiments (N=2). Results (mean ± SEM) are given as percentage of control (APP+/+). Scale bar: 200µm. *p=0.0220, Mann-Whitney test. **D)** Neurons harvested at DIV7 and cell extracts analyzed by Western blotting for GABARα1 and GADPH expression. Quantification of GABARα1 was normalized to GAPDH expression. Results (mean ± SEM) are expressed as percentage of Ct (n=5, N=2). *p=0.0197, Student’s t-test.

Finally, we evaluated whether impairments affecting GABAergic neurotransmission components we observed in vitro *in vitro* could are relevant in the brain. We quantified the expression of GAD65 in cortices and hippocampi of 3 month old mice deficient for APP (APP-/-) compared to their wild-type counterpart (APP+/+). Consistent increase in GAD65 expression is observed both in cortex (Figure 6A) and in hippocampus (Figure 6B) of APP-/- mice; indicating that GABAergic neurotransmission component GAD65 is also affected in adult mouse brain and supporting our *in vitro* findings.

**Figure 6:**
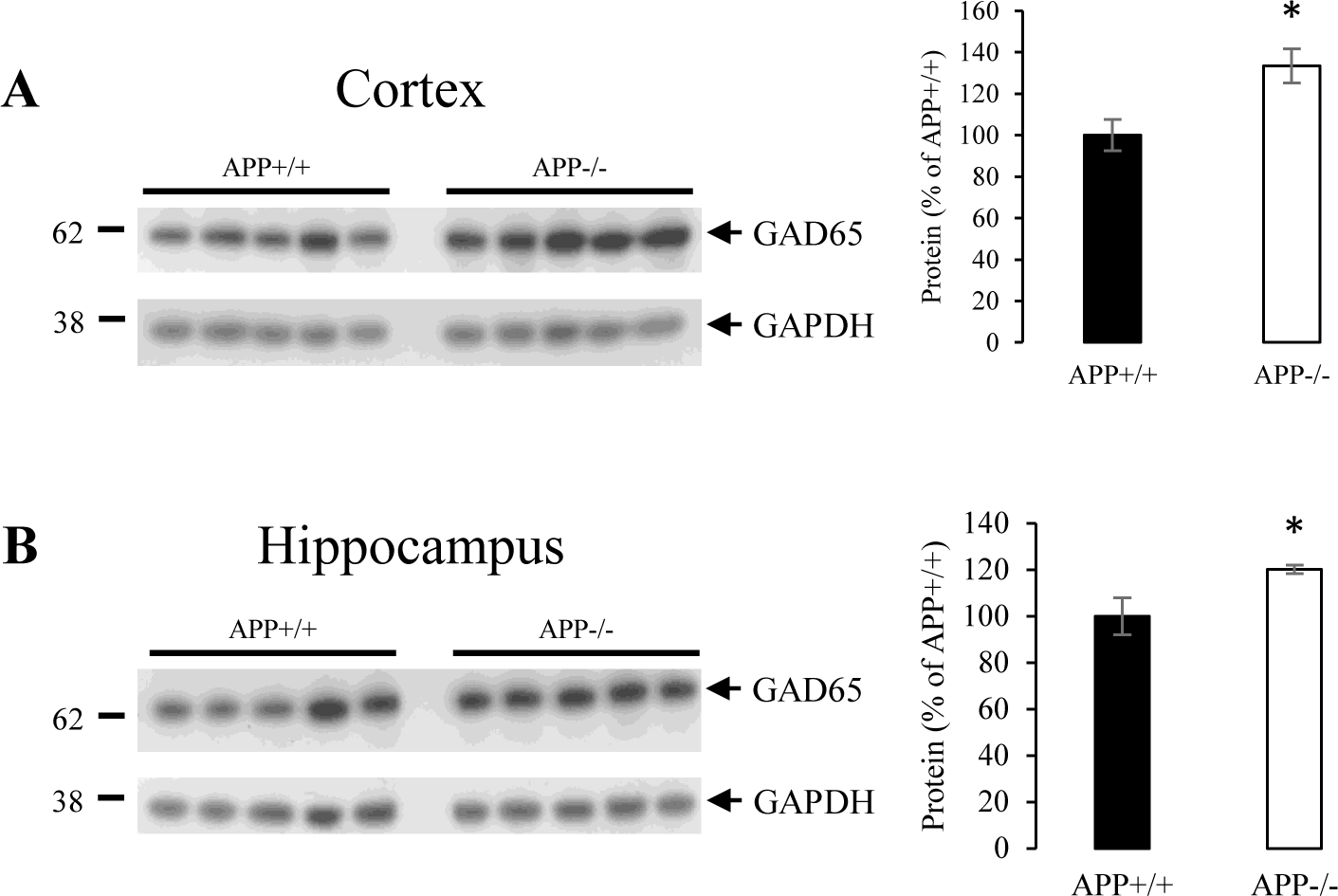
**GAD65 expression in cortex and hippocampus of adult mice A**) *Left panel* Western blot analysis of GAD65 and GAPDH expression in cortex of 3 month old APP+/+ and APP-/- mice (N=5). *Right panel* Quantification of GAD65 was normalized to GAPDH expression. Results (mean ± SEM) are expressed as percentage of APP+/+ (N=5). *p=0.0166, Student’s t-test. **B**) *Left panel* Western blot analysis of GAD65 and GAPDH expression in hippocampus of 3 month old APP+/+ and APP-/- mice (N=5). *Right panel* Quantification of GAD65 was normalized to GAPDH expression. Results (mean ± SEM) are expressed as percentage of APP+/+ (N=5). *p=0.0404, Student’s t-test.

### Phenotype of NPAS4-deficient neurons mimics APP deficiency

As for APP, we used the CRISPR-Cas9 approach in order to silence *Npas4* and analyze whether NPAS4 deficiency could recapitulate a major trait observed in APP-/- neurons, i.e. imbalance of inhibitory transmission by the upregulation of GABA release. Given that (i) CRISPR-Cas9 editing is hard to evaluate by quantifying mRNAs expression, (ii) that available antibodies poorly detect NPAS4 in basal conditions, we decided to check the down-regulation of *Npas4* gene expression by measuring NPAS4 protein upon depolarization by KCl (Lin et al., 2008). CRISPR-Cas9-induced silencing resulted in a decrease in NPAS4 by approximately 50% (Figure 7A), a similar extent to that observed of mRNAs in APP-/- neurons at DIV7 (Figure 1D). This downregulation of NPAS4 is not due to a lentiviral toxic effect (Figure S4C). Strikingly, like for APP-deficient neurons (Figure 5C), NPAS4-deficient neurons showed an increase in GAD65 staining (Figure 7B-C), GAD65 protein expression (Figure 7D), and GABA release in the medium (Figure 67C) when compared to control neurons. We measured the expression of GABA receptor subunit alpha 1 (GABARa1) at DIV7 and observed, like in APP-/- primary neurons, a decrease in protein expression after *Npas4* knockdown (Figure 7D).

**Figure 7:**
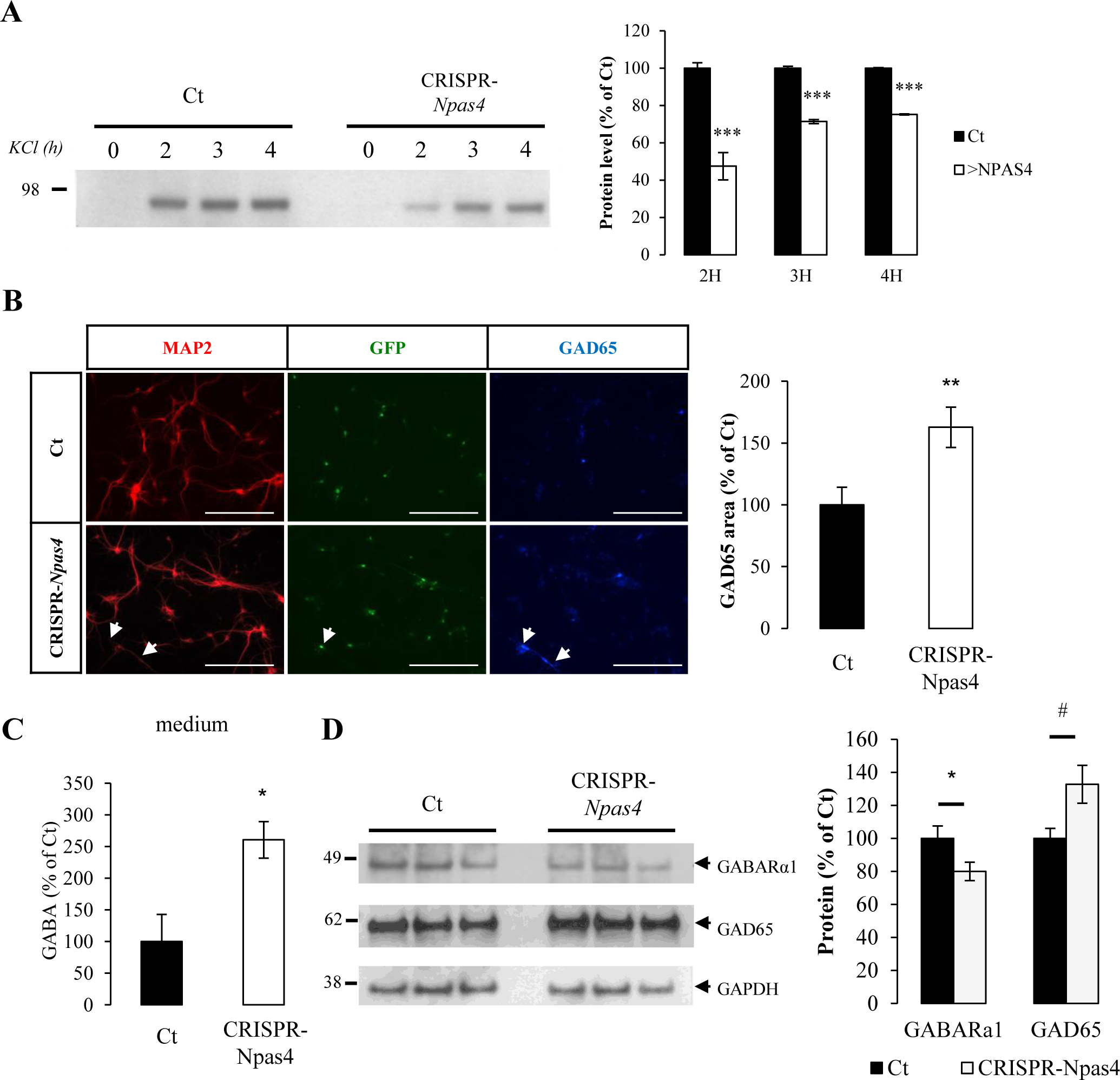
**Npas4 silencing by CRISPR-Cas9 mimicks cell phenotype observed in APP deficient neurons. Changes on inhibitory (GABA) synapses was analyzed after *Npas4* silencing A**) *Left panel.* Cortical neurons infected with CRISPR-Cas9 lentivirus targeting *Npas4* gene (CRISPR-Npas4) show reduced NPAS4 levels as measured by Western blotting after membrane depolarization with 50mM potassium chloride (KCl). Control viruses without sgRNA were used as controls (Ct). Note that NPAS4 is barely detectable in non-depolarized neurons (Ct). *Right panel.* Quantification of NPAS4 protein level after 2, 3 and 4h of KCl depolarization. Results (mean ± SEM) are expressed as percentage of non-treated controls Ct (N=2). ***p<0,0001 Student’s t-test. **B**) Cortical neurons infected with CRISPR-Npas4 lentiviruses at DIV1 were immunostained against MAP2 and glutamate decarboxylase 65 (GAD65) at DIV7. Quantification of GAD65 signal was normalized to the number of cells (5 fields per coverslip, 2 coverslips for each genotype in two independent experiments, N=2). Results (mean ± SEM) are given as percentage of control (Ct). Scale bar: 200µm. **p=0.0024. Mann-Whitney test. **C**) Quantification of γ-amino butyric acid (GABA) in culture medium at DIV7of infected control neurons (Ct) and CRISPR-*Npas4* infected neurons. Results (mean ± SEM) are expressed as percentage of Ct (n=5, N=2). *p=0,0146, Student-t test. **D**) Neurons harvested at DIV7 and cell extracts analyzed by Western blotting for GABARα1, GAD65 and GADPH expression. Quantification of GABARα1 and GAD65 were normalized to GAPDH expression. Results (mean ± SEM) are expressed as percentage of Ct (n=8, N=3). *p=0.049, ^#^p=0.0247, Student’s t-test.

## Discussion

One major APP function is to control synaptic formation, transmission and plasticity (Muller et al., 2017). We showed here that APP deficiency in cortical neurons impairs the balance between excitatory and inhibitory synaptic markers, and that this process relies on the activity-dependent transcription factor NPAS4. We initially identified the *Npas4* IEG as a potential APP target gene by a non-biased transcriptome profiling approach. The APP-dependent regulation of *Npas4* expression involves its extracellular domain (sAPPα) but not AICD. APP appears to exert a fine tuning of excitatory/inhibitory synaptic inputs in neurons and its absence enhances, through the downregulation of *Npas4*, inhibitory GABAergic transmission.

### APP-dependent expression of *Npas4* in differentiated neuronal culture

The transcriptome analysis of APP+/+ vs. APP-/- neurons at embryonic day 17 (E18-DIV0) and at different stages of primary cortical neuron differentiation (DIV3-DIV7) indicated that the transcriptional changes in the absence of APP were moderate. This unexpected result is however in line with a comparative transcriptome study of APP family members in the adult mouse cortex (Aydin et al., 2011). One possible explanation to the subtle effects of APP deficiency on the transcriptome can be functional compensation by APLPs (Shariati and De Strooper, 2013). However, we did not measure any changes in APLP1 and APLP2 expression in our APP-/- models in agreement with previous observations total brain extracts (Zheng et al., 1995) or in primary cortical neurons (White et al., 1998). Transcriptional modifications we measured are thus related to APP *per se*. APP-dependent transcriptional regulations are subtle, and likely to act by fine-tuning classes of gene involved in neuronal pathway rather than single target genes. In addition, we found that none of the APP/AICD target genes were differentially expressed in APP-/- neurons at DIV3-DIV7 or at E18 (See Supplementary Table 3). The identification of AICD-dependent gene expression stemmed from studies carried out in an array of *in vitro-* and -to a lesser extent- *in vivo* models (for review see Pardossi-Piquard and Checler, 2012; Grimm et al., 2013). Some of these findings were confirmed or debated in subsequent investigations (Hebert et al., 2006; Chen and Selkoe, 2007; Waldron et al., 2008; Aydin et al., 2011). AICD-dependent gene transcription and how it relates to APP function appears thus, if not controversial, scarcely understood.

We found that the expression of *Npas4*, an activity-dependent IEG, is downregulated in the absence of APP and particularly at DIV7. *Npas4* downregulation was observed it in APP-/- primary neurons and upon acute APP knock-down by a CRISPR-Cas9 approach (Figure 3D), establishing a causal relation between APP and *Npas4* transcription. Furthermore, APP-dependent *Npas4* expression at DIV7 does not rely on AICD release, although DIV7 corresponds to the differentiation stage where AICD is readily produced by neurons (Kimberly et al., 2005). Previous studies indicated that regulation of some APP target genes does not require the generation of AICD (Hicks et al., 2013). Quite strikingly, DAPT treatments, used to block AICD production in our setup, increased Npas4 expression in APP+/+ neurons. This effect might imply that inhibition of γ-secretase increases the neuronal activity in an APP-dependent manner (APP-/- neurons showed no modification of *Npas4* expression after treatment). In line with this, γ-secretase inhibition was shown to increase excitatory postsynaptic currents (EPSCs) (Priller et al., 2006; Restituito et al., 2011). Further investigations are required to understand this observation. However, one hypothesis could be that the loss of Aβ underlies the DAPT effects we observed. Several studies reported that Aβ depresses AMPA- and NMDA-receptor mediated currents and EPSCs in neurons arguing toward a negative feedback of Aβ on synaptic transmission (Kamenetz et al., 2003; Snyder et al., 2005; Hsieh et al., 2006). This feedback is not possible in a APP-/- background. Alternatively, studies indicated that inhibition of γ-secretase induces an increase of production of sAPPα (Chen et al., 2015). In that case, increased activity related to increased sAPPα production would corroborate the results we obtained on *Npas4* expression by treating neurons with sAPPα. We also found that sAPPα effects on Npas4 expression are observed only in APP+/+ and not in APP-/- background. This observation indicates that (i) the transcriptional effects of sAPPα require the presence of endogenous APP holoproteins (ii) homophilic ectodomain interactions are likely to be involved. Soluble APP has been shown to rescue many traits of APP-deficient mice (Ring et al., 2007; Weyer et al., 2014) and was suggested to promote its physiological effects by interaction with APP holoprotein (Milosch et al., 2014; Deyts et al., 2016).

### Alteration of GABAergic inputs in APP deficient neurons are related to Npas4 downregulation

In the absence of APP, we observed an increase in neuronal outgrowth and GAD65 signal, as well as increased GABA release in the medium. To note, *Npas4* knockdown mimics APP deficiency on GAD65 levels and GABA measurements (Figure 8). This supports the hypothesis APP regulates the fine-tuning inhibitory synaptic transmission in the neuronal network through NPAS4. First, this is in agreement with very recent work showing that APP regulates GABAergic neurotransmission during neuronal differentiation (Doshina et al., 2017). *In vivo* studies evidenced increased GABA levels in the brain of APP-/- mice (Lee et al., 2010). Secondly, the finding that APP-dependent neuronal processes are mediated by NPAS4 is relevant to experimental evidences reported in previous studies. NPAS4 possesses unique features among the IEGs (Sun and Lin, 2016): (i) it is only expressed in neurons; (ii) it is activated selectively by neuronal activity; (iii) it has been shown to be important to shape glutamatergic and GABAergic synaptic inputs. NPAS4 is implicated in a transcriptional program that regulates neuronal firing responses to excitatory transmission by enhancing inhibition (Lin et al., 2008), and is critical for the homeostatic mechanisms that keep neuronal firing in response to stimuli within normal levels (Spiegel et al., 2014). Increasing the excitability of a set of neurons leads to changes in both their input and axonal synapses. NPAS4 is necessary for modulating the inputs synapses but not the axonal synapses of these neurons (Sim et al., 2013). NPAS4 is induced in excitatory neurons, where it promotes increased numbers of inhibitory synaptic inputs. (Spiegel et al., 2014). Altogether, these specific functions of NPAS4 correlate well with our main observation that APP-dependent *Npas4* expression is related to the upregulation of the GABAergic system in APP-deficient neurons. This is not restricted to primary neurons, since we measured an increase in markers of GABAergic synapses in adult mouse brain.

The overall effect of APP deficiency of neuronal network activity and synaptic transmission needs further neurophysiological investigations that are beyond the scope of the present study. Important points must be kept in mind here. First, GABAergic transmission shifts from excitatory to inhibitory during development (Ben-Ari, 2002), and our findings should be evaluated by electrophysiological recordings in mature neurons. For instance, we found that the level of GABA receptor subunit alpha 1 (GABARα1) was diminished in the absence of APP and in NPAS4-deficient neurons. This is in agreement with recent study showing that GABARα1 is particularly decrease in hippocampus of APP-/- mice (Chen et al. 2017) correlating with a decrease in IPSC amplitude. But on the other hand, it suggests that increases in GABA release in APP-deficient models may not result in a net increase of inhibitory transmission, or at least there is a complex modulation of neuronal response to GABA.

### Possible relevance to the AD pathophysiology

APP plays a central role in the onset and progression of AD by releasing the Aβ peptide, but, APP deficiency is more difficult to correlate to the pathology. Still, it is admitted that impairment of APP function *per se*, either caused by FAD mutations or upon ageing, may contribute to neuronal dysfunction occurring in the disease. For instance, the phenotype of APP deletion in the CNS is age-dependent (Priller et al., 2006). Upon aging, impairments in learning and memory associated with deficits in LTP are observed in APP-deficient mice (Ring et al., 2007). The role of APP in maintaining spine architecture is supported by the reduction in dendritic length and branching as well as in total spine density in old APP-deficient mice (Lee et al., 2010; Tyan et al., 2012). A severe decrease in metabolic activity was also observed in presynaptic densities of APP KO animals (Lassek et al., 2017). This is an important feature, because bioenergetics and metabolic activity are fueling the synthesis of neuromediators (glutamate and consequently GABA), and providing energy supply and calcium buffering essential for synaptic function and plasticity.

Significantly lower levels of GABA and glutamate were measured in the temporal cortex of AD patients, pointing to deficient synaptic function and an imbalance in neuronal excitatory/inhibitory transmission (Gueli and Taibi, 2013). These observations unambiguously support changes in neurotransmission in AD (and even in ageing brain), but the mechanisms underlying this process are hardly understood. Here we found that neuronal activity by itself, sensed by the NPAS4 IEG, reshapes synaptic GABAergic inputs on neurons, in line with recently reported modifications of GABA transmission in AD models (Doshina et al., 2017). Very interestingly, NPAS4 expression decreases along with AD progression, particularly at Braak NFT stages (I-II) corresponding to lesions developed in transentorhinal/entorhinal cortex (Miyashita et al., 2014). Downregulation of GABAergic transmission could also underlie the increased risk for unprovoked seizures observed in individuals with AD compared to non-demented individuals of the same age (Friedman et al., 2012).

In conclusion, our main observation that APP deficiency in neurons is integrated by the activity-dependent NPAS4 IEG to further re-modulate inhibitory and excitatory neuronal inputs, provides new insight to understand the role of APP in synaptic activity, but also a mechanistic frame to further explore the impairments of network activity in AD.

**Abbreviations**
Aβ: amyloid-β peptide
AChE: acetylcholinesterase
AD: Alzheimer’s disease
AICD: APP intracellular domain
APP: Amyloid Precusor Protein
APLP1: APP-like Protein1
APLP2: APP- like Protein 2
PPD: paired pulse depression
CRISPR: clustered regularly interspaced short palindromic repeats
DAPT: N-[N-(3,5-Difluorophenacetyl)-L-alanyl]-S-phenylglycine t-butyl ester
ECM: extracellular matrix
GABARa1: GABA(A) receptor subunit alpha-1
GAD65: glutamate decarboxylase 65
GRIK1: glutamate ionotropic receptor kainate type subunit-1
IEG: immediate early gene
KA: kainic acid
LTP: long term potentiation
NPAS4: neuronal PAS domain protein 4
sAPPα: soluble APP alpha

## Acknowlegments

pLenti CMV/TO Puro empty (w175-1) was a gift from Eric Campeau & Paul Kaufman (Addgene plasmid #17482). pL-CRISPR.EFS.GFP (Addgene plasmid # 57818) and pL-CRISPR.EFS.tRFP (Addgene plasmid # 57819) were a gift from Benjamin Ebert. pCMV delta R8.2 (Addgene plasmid # 12263) and pMD2.G (Addgene plasmid # 12259) were a gift from Didier Trono. We thank Jerome Ambroise for insight and technical support in the analysis of microarray data. We thank Nicolas Dauguet for the cell cytometry sorting of the neurons. We thank Devkee Mahesh Vadukul for her critical and linguistic revision of the manuscript.

## Author contributions

R.O. and P.K.C. designed the research study; R.O. conducted the experiments, with the help of B.T., S.C., C.V., A.D., N.P. and F.P. Intracellular calcium measurement were designed and performed with the help of P.D. and M.V. All the authors analyzed data. R.O. and P.K.C. wrote the manuscript with the inputs of S.S., N.P. and J.N.O. All the authors have read and approved the final manuscript.

---

**Figure S1:**
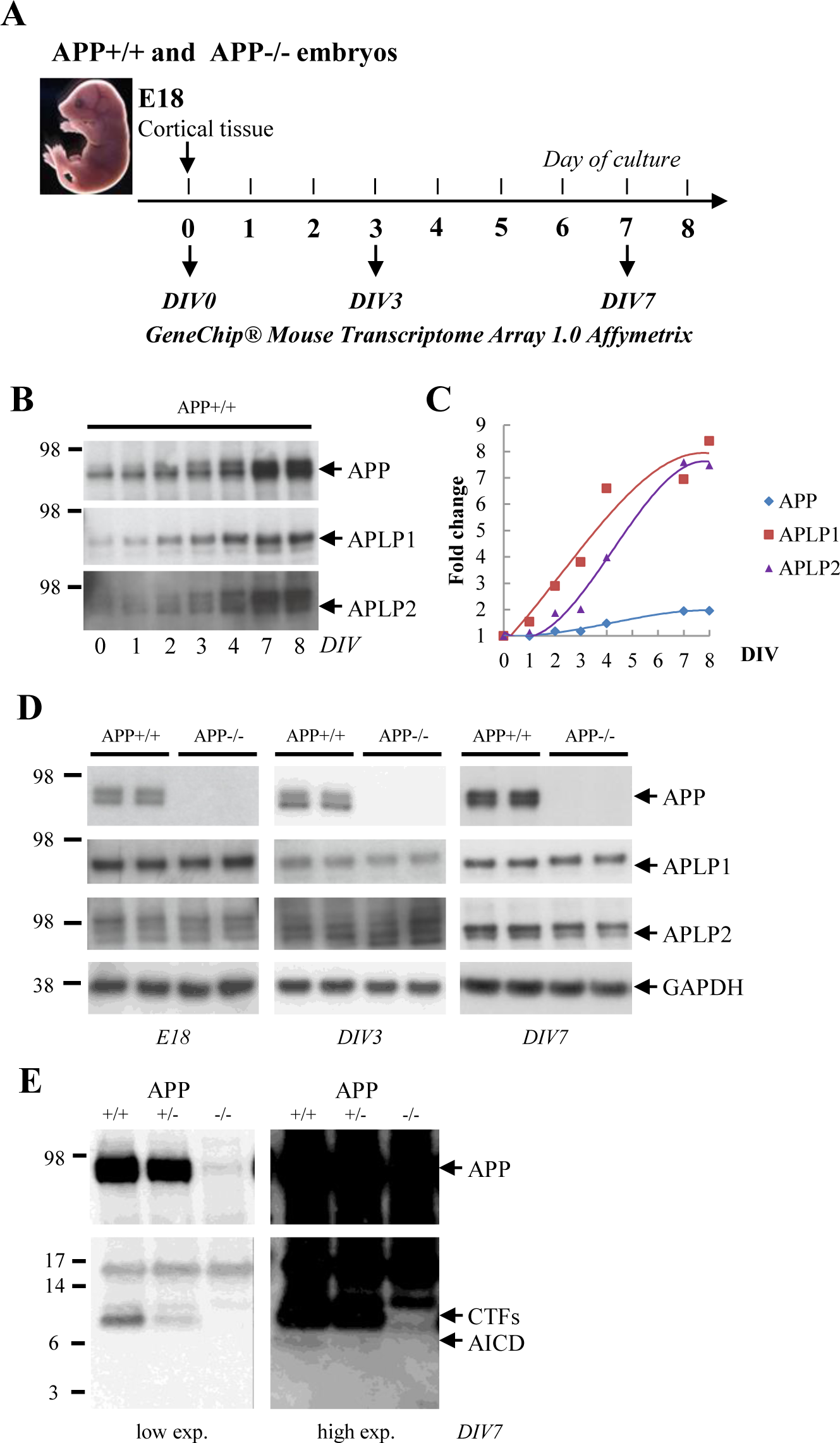
**Experimental workflow and model characterization A**) Experimental design used for the study. Cortical tissue was taken at embryonic day 18 (E18); neurons were cultured and experiments were mainly carried out after 3 and 7 days in vitro (DIV3 and DIV7). Transcriptome analysis was performed on embryonic cortex (E18) and at DIV3 or DIV7. **B**) APP, APLP1 and APLP2 expressions were analyzed by Western blotting at the indicated days of culture in APP+/+ neurons. **C**) Quantification of APP, APLP1 and APLP2 expression over time in APP+/+ neurons. Accumulation is represented as fold change over the signal measured at day 0. Quantification was performed from one neuronal culture **D**) APLP1 and APLP2 expressions are not modified in cortical tissue at E18 and primary neuron cultures at DIV3 and DIV 7. Expression of APP, APLP1, APLP2 was analyzed by Western blotting of cells lysates from APP+/+ and APP-/- primary neuron cultures. **E**) Samples from primary cultures at DIV7 (APP+/+, APP+/- and APP-/- neurons) were probed (Western blotting) with an antibody directed against APP C-terminus for APP C-terminal fragments (CTFs) and AICD. Low and high exposures of a typical blot are shown. Arrows indicate the expected position of APP holoprotein, APP CTFs and AICD.

**Figure S2:**
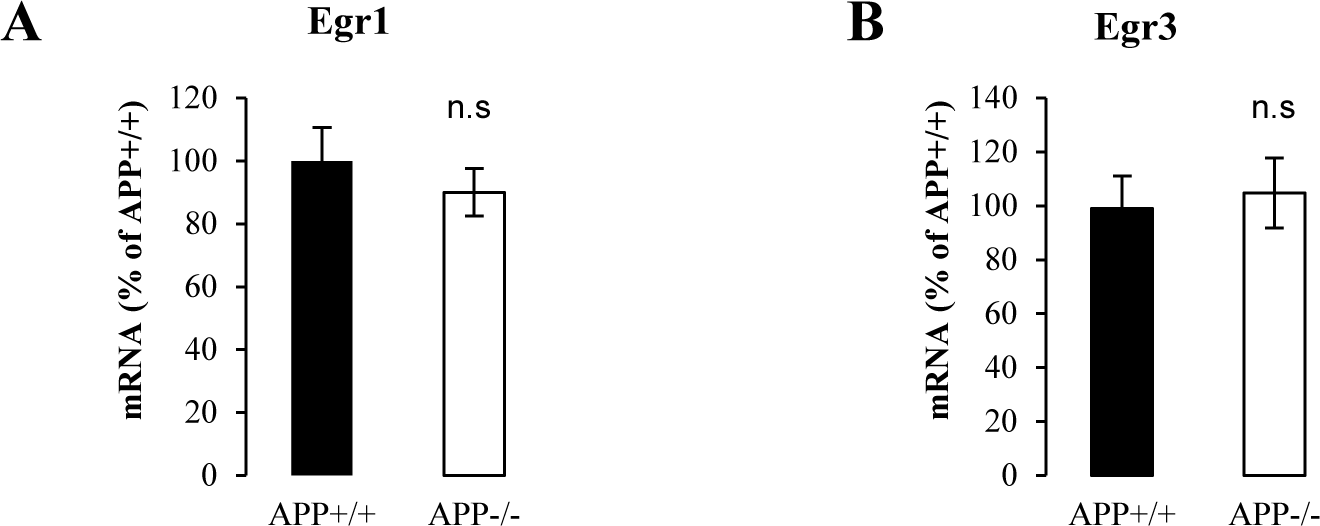
**Egr1 and Egr3 expressions are not modified in APP deficient neurons** Egr1 and Egr3 expressions were evaluated in APP+/+ vs. APP-/- primary neurons at DIV7. **A**) Egr1 mRNA level was measured by qPCR (n=6, N=3) at DIV7. Results (mean ± SEM) are given as percentage of controls (APP+/+) n.s= non-significant, Student’s t-test. **B**) Egr3 mRNA level was measured by qPCR (n=6, N=3) at DIV7. Results (mean ± SEM) are given as percentage of controls (APP+/+) n.s= non-significant, Student’s t-test.

**Figure S3:**
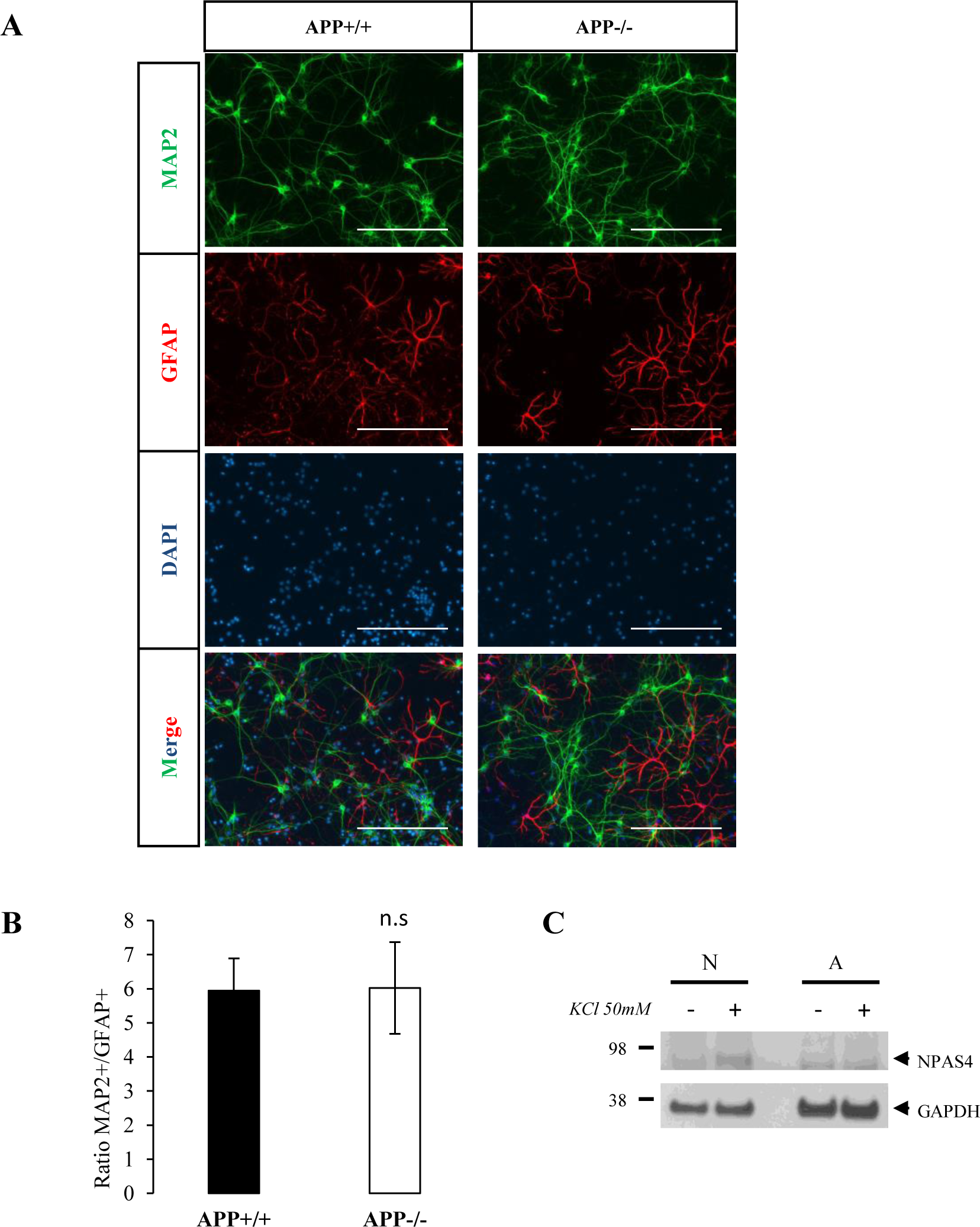
**Astrocytes in primary neuron culture and their implication in Npas4 expression. A**) Primary culture of cortical neurons at DIV7. Cultures were immunostained with the neuron specific protein MAP2 (green), the glial specific protein GFAP (red) and the DAPI (light blue). Scale bar = 400µm. **B**) Quantification of neurons (MAP2+) and astrocytes (GFAP+) in the primary cortical culture. At least five fields per coverslip were analyzed for APP+/+ and APP-/- cultures in two independent experiments (n≥5, N=2). Results are expressed as the ratio of MAP2+ (neurons) and GFAP+ (astrocytes) (mean ± s.e.m). n.s= non-significant, Mann-Whitney test. **C**) Western blotting analysis of Npas4 induction in neurons (N) and astrocytes (A) after depolarization with 50mM potassium chloride (KCl) for 2 hours.

**Figure S4:**
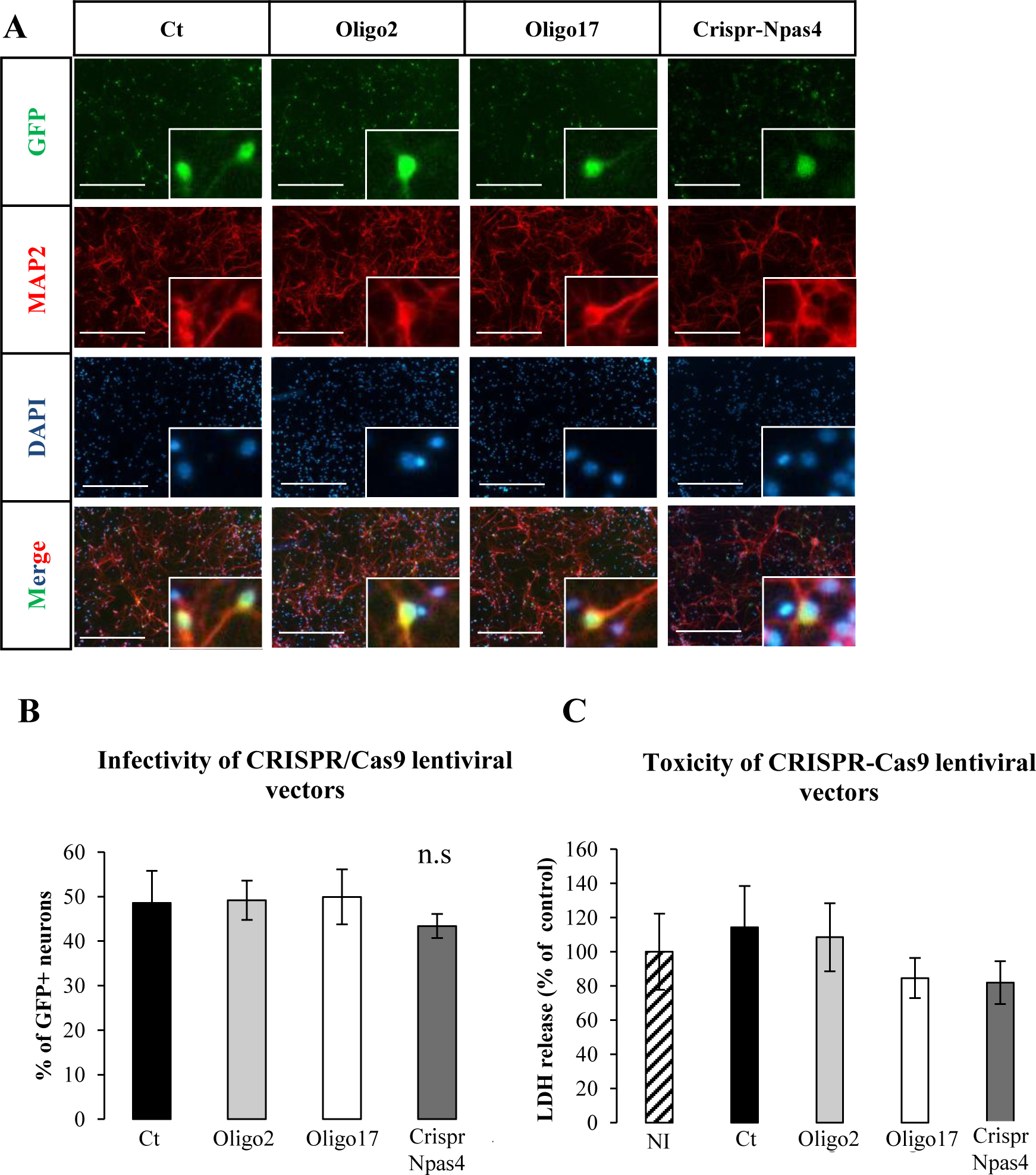
**Infectivity and toxicity of lentiviral CRISPR-Cas9 vectors A**) Cortical neurons were infected at DIV1 with lentiviruses expressing sgRNAs (Oligo2, Oligo17 or CRISPR-*Npas4*) or no sgRNA (Ct), SpCas9 and GFP. Cultures were immunostained for MAP2 (red) and DAPI (light blue) at DIV7. Scale bar = 400µm. **B**) Quantification of GFP+ neurons (GFP+/MAP2+) in total neuron population (MAP2+) after lentiviral CRISPR-Cas9 infection with control (Ct), Oligo2, Oligo17 or CRISPR-*Npas4*. At least five fields were analyzed for each lentiviral vector in two independent experiments (n≥5, N=2). Results are expressed as percentage of GFP+/MAP2+ cells in total MAP2+ cells (mean ± s.e.m). n.s= non-significant, Kruskal-Wallis test and Dunn’s multiple comparison test. **C**) Measurement of LDH activity released after infection (DIV7) of primary neuron with control (Ct), Oligo2, Oligo17 or CRISPR-*Npas4* at DIV7 lentiviral vectors. Background LDH release was determined in non-infected control cultures (NI). Results were expressed as percentage of total LDH release measured in non-infected control cultures (NI) in 2 independent experiments (n=12, N=2).

**Figure S6:**
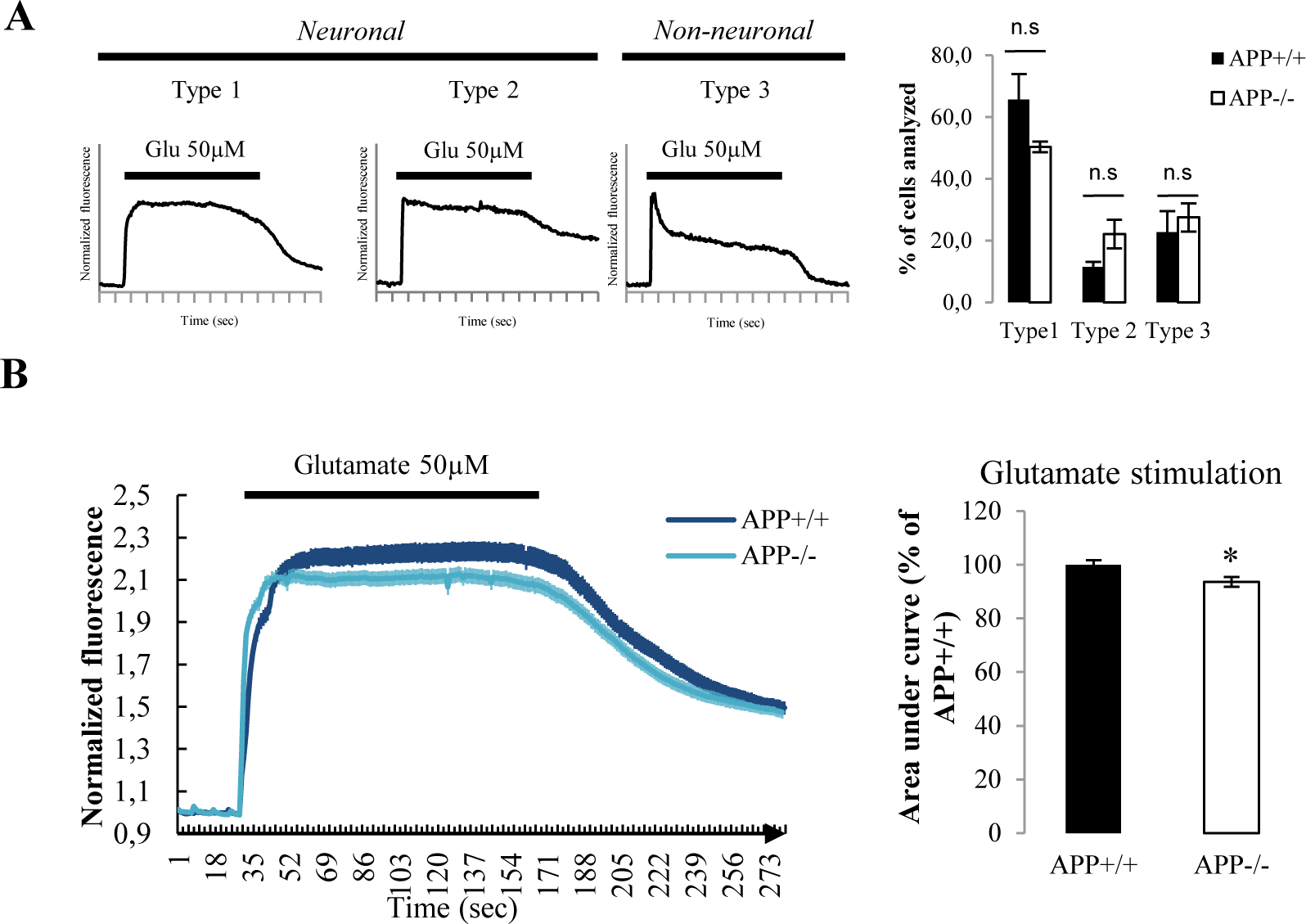
**Modification in glutamate responses in APP-/- neurons measured by intracellular calcium imaging.** Neuronal activity was measured at DIV7 by calcium imaging. A) Left panel. Different calcium responses were observed after stimulation with 50 µM glutamate and classified as described by Prickering and co-workers (Prickering et al. 2008) between neuronal and non-neuronal responses. To note X-axe graduation correspond to 20 sec. Right panel. The proportion of cells displaying Type 1, 2 or 3 response was quantified in three independent experiments (n=9, N=3). n.s.= non-significant. Student-t test. B) Normalized fluorescence trace (mean ± SEM) measured in APP+/+ and APP-/- neurons upon perfusion for 150 sec with 50 µM glutamate. The area under curve (AUC) was quantified for 50 neurons per coverslips. A total of 9 coverslips for each genotype was recorded in three independent experiments (N=3). The graph on the right shows AUC expressed as percentage of control (APP+/+). *p=0,0106, Student’s t-test.

**Figure S6:**
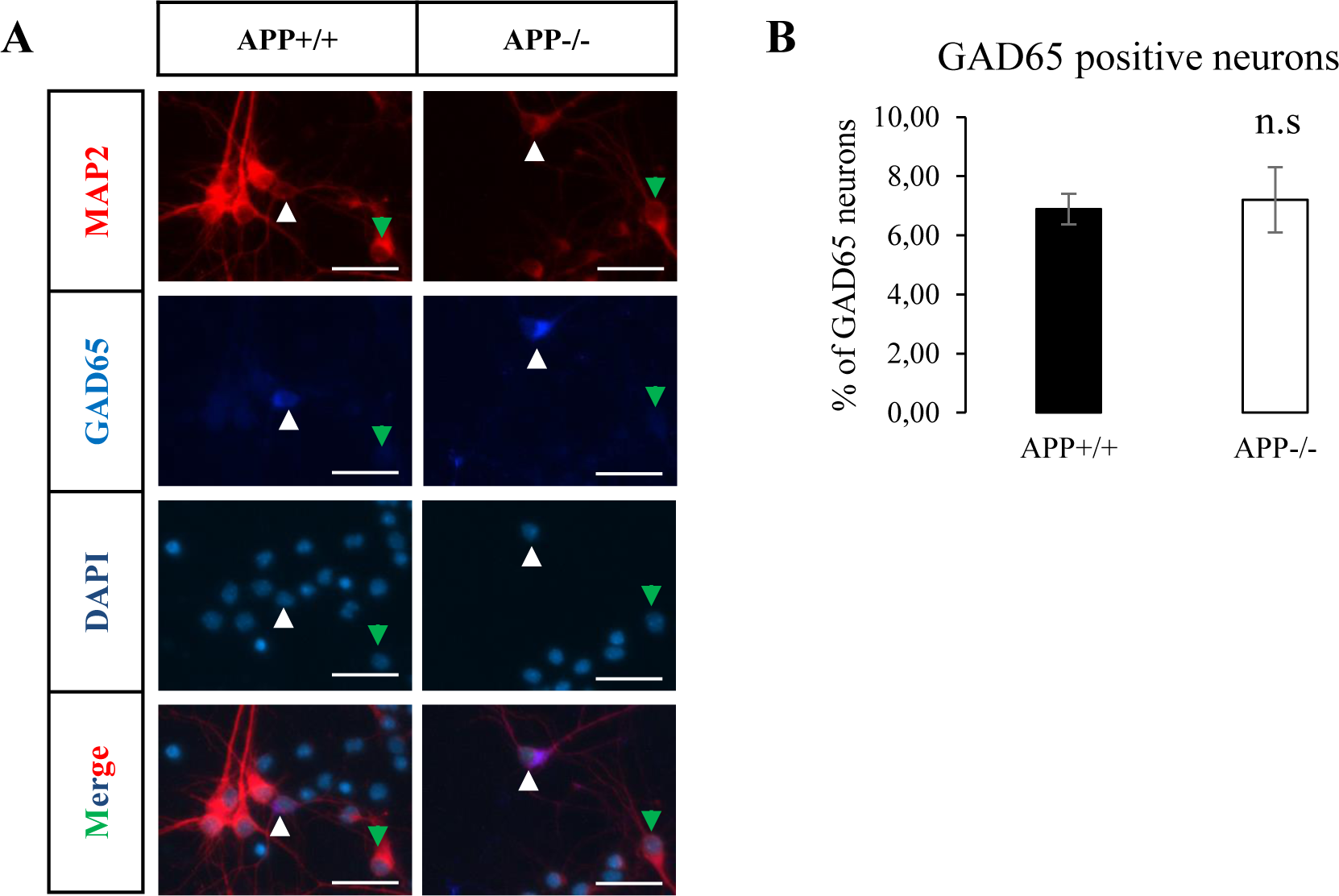
**GAD65 positive neurons in primary cortical culture.** A) Primary culture of cortical neurons after at DIV7. Cultures were immunostained with the neuron specific protein MAP2 (red), GAD65 (dark blue) and DAPI (light blue). Representative 20x micrographs show GAD65 positive neurons (white arrowhead) and GAD65 negative neuron (green arrowhead). **B**) Images (20x objective) were quantified (10 fields per coverslip for each genotype) in three independent cultures (n=30, N=3). Results (mean ± s.e.m) are expressed as percentage of GAD65+ MAP2+ cells (GAD65+ neurons) among all MAP2+ cells (neurons). n.s= non-significant, Mann-Whitney test. Scale bar = 20µm.

