## Supplemental tables for "Amyloid Precursor Protein (APP) controls excitatory/inhibitory synaptic inputs by regulating the transcriptional activator Neuronal PAS Domain Protein 4 (NPAS4)"

### 1 Supplementary Material

#### 1.1 Supplementary Table 1: qPCR primers

| **Primer** | **Forward** | **Reverse** |
|---|---|---|
| *Gapdh* | 5’-ACCCAGAAGACTGTGGATGG-3’ | 5’- ACACATTGGGGGTAGGAACA-3’ |
| *Npas4* | 5’-GCTATACTCAGAAGGTCCAGAAGGC-3’ | 5’-TCAGAGAATGAGGGTAGCACAGC-3’ |
| *Egr1* | 5’-TCCTCTCCATCACATGCCTG-3’ | 5’-CACTCTGACACATGCTCCAG-3’ |
| *Egr3* | 5’-GACTCGGTAGCCCATTACAATC-3’ | 5’-ACTTTCCCAAGTAGGTCACGG-3’ |
| *Grik1* | 5’-GCTGGACACTCCTCTATTTCCT-3’ | 5’-ACAGGTTCGTTTTCCACAGTT-3’ |

**Supplementary Table 1: qPCR primers**

Sequences of forward and reverse primers for quantitative PCR for genes analyzed in the study.

#### 1.2 Supplementary Table 2: CRISPR-Cas9 sgRNA

| gRNA | Sequence | PAM – Specificity score |
|---|---|---|
| Oligo2 | GTGGAAGATCCGCCGCGCCC | TGG - 95 |
| Oligo17 | GTACCCACTGATGGCAACGC | CGG - 92 |
| CRISPR-*Npas4* | GACCCTTGCGAGTGTAGATGC | AGG - 83 |

**Supplemental Table 2: CRISPR-Cas9 sgRNA**

Sequences, PAM motifs and scores for each sgRNAs used in the study.

#### 1.3 Supplementary Table 3: AICD dependent target genes are not differentially expressed in differentiating neuronal culture

| AICD-target genes | | | Results in our array | | |
|---|---|---|---|---|---|
| **Gene symbol** | ***References*** | **Up or down** | **E18** | **3DIV** | **7DIV** |
| **APP** | *von Rotz et al. 2004* | Up | 0,07 | 0,07 | 0,04 |
| **RA-responsive genes** | *Gao and Pimplikar 2001* | Down | - | - | - |
| **KAI1/CD82** | *Baek et al. 2002; von Rotzet al. 2004* | Up | 1,07 | 0,97 | 0,97 |
| **GSK3b** | *Kim et al. 2003; von Rotzet al. 2004;* | Up | 0,99 | 0,99 | 1,04 |
| **BACE** | *von Rotz et al. 2004* | Up | 0,98 | 1,06 | 1,04 |
| **Tip60** | *von Rotz et al. 2004* | Up | 1,03 | 1,01 | 0,99 |
| **NEP** | *Pardossi-Piquard et al.2005; Belyaev* | Up | 0,97 | 1,01 | 0,99 |
| **p53** | *Alves da Costa et al. 2006 Ozaki et al. 2006* | Up | 0,99 | 1,04 | 1,07 |
| **a2-Actin** | *Muller et al. 2007* | Up | 1,02 | 1,09 | 0,92 |
| **Transgelin** | *Muller et al. 2007* | Up | 1,01 | 0,97 | 0,97 |
| **IGFBP3** | *Muller et al. 2007* | Up | 1,02 | 1,02 | 0,97 |
| **EGFR** | *Zhang et al. 2007* | Down | 0,93 | 1,05 | 0,93 |
| **LRP1** | *Liu et al. 2007* | Down | 1 | 1,07 | 1,04 |
| **Cyclin B1 and D1** | *Ahn et al. 2008* | / | / | / | / |
| **VGLUT2** | *Schrenk-Siemens et al. 2008* | Up | 0,96 | 1,05 | 1,23 |
| **CHOP** | *Takahashi et al. 2009* | Up | 1 | 1,04 | 0,93 |
| **Aquaporin 1** | *Huysseune et al. 2009* | Up | 1,44 | 0,95 | 0,96 |
| **S100a9** | *Ha et al. 2010b* | Up | 1,15 | 0,92 | 0,99 |

| | | | | | |
|---|---|---|---|---|---|
| **ApoJ/clusterin** | *Kogel et al. 2011* | Down | 1,16 | 1,12 | 1,02 |
| **Ptch1** | *Trazzi et al. 2011* | Up | 1,03 | 1,06 | 1,01 |

**Supplementary Table 3: AICD dependent target genes are not differentially expressed in differentiating neuronal culture**

Summary of the transcriptome analysis for the AICD target genes reported to be transcriptionally regulated by AICD (adapted from Pardossi-Piquard and Checler, 2012). First column correspond to Gene Symbol. Second column shows to the associated publication of the corresponding gene and the third column correspond to the direction of regulation by the AICD (Up or down regulated). The last three columns depicted the linear fold change (APP-/- vs APP+/+) observed for each gene under our experimental conditions E18 (DIV0), DIV3 and DIV7. APP fold change is highlighted in red and taken as a positive control.
